# ATM and 53BP1 regulate alternative end joining-mediated V(D)J recombination

**DOI:** 10.1101/2024.04.25.591195

**Authors:** Jinglong Wang, Cheyenne A. Sadeghi, Long V. Le, Marie Le Bouteiller, Richard L. Frock

## Abstract

G0/G1-phase alternative end joining (A-EJ) is a recently defined mutagenic pathway characterized by resected deletion and translocation joints that are predominantly direct and are distinguished from A-EJ in cycling cells which rely much more on microhomology-mediated end joining (MMEJ). Using chemical and genetic approaches, we systematically evaluate potential A-EJ factors and DNA damage response (DDR) genes to support this mechanism by mapping the repair fates of RAG1/2-initiated DSBs in the context of Igκ locus V-J recombination and chromosome translocation. Our findings highlight a polymerase theta-independent Parp1-XRCC1/Lig3 axis as central A-EJ components, supported by 53BP1 in the context of an ATM-activated DDR. Mechanistically, we demonstrate varied changes in short-range resection, MMEJ, and translocation, imposed by compromising specific DDR activities, which include polymerase alpha, ATR, DNA2, and Mre11. This study advances our understanding of DNA damage repair within the 53BP1 regulatory domain and the RAG1/2 post-cleavage complex.

## Introduction

V(D)J recombination assembles the variable region of antigen receptor loci in the G0/G1 phase of the cell cycle and is essential for B and T cell development. Recombination is highly-coordinated, involving the loading of the RAG1/2 endonuclease at J region recombination centers, pairing of D or V gene segments across chromatin loops, RAG1/2 incision, hairpin opening and processing of the coding ends, ligation of coding ends and, separately, ligation of blunt recombination sequence (signal) ends (1, 2). The primary mechanism for ligation is nonhomologous end joining (NHEJ). However, core NHEJ deficiency (i.e. Ku70, Ku80, XRCC4, Lig4) reveals an alternative end joining (A-EJ) machinery that completes DNA double strand break (DSB) repair to varying degrees. For instance, while A-EJ in the absence of XRCC4 or Lig4 deficiency is extremely low in G0/G1 phase, it is quite robust in cycling phases, characterized by kilobase-long resection and near exclusive junctional microhomology (MH) utilization (3, 4). In contrast, G0/G1 A-EJ in the absence of Ku70 is robust, though less efficient than NHEJ, repairing both Cas9 and RAG1/2 DSBs with a limited resection window and a greater direct to MH ratio that more closely resembles NHEJ than *Lig4^-/-^*(4). However, a key discerning feature for V(D)J recombination by either A-EJ mechanism is the significant loss in biased joining of coding ends to each other and likewise for signal ends. Thus, end joining in the absence of Ku70/80 represents a true A-EJ mechanism; whereas the absence of XRCC4/Lig4, represents an end joining mechanism influenced by NHEJ intermediates (4).

Despite its significance in tumorigenesis (5), A-EJ mechanisms remain poorly understood. Early identified components in cycling cells included Parp1, XRCC1, and Lig3 (6–7), which contribute to chromosome translocation in mouse embryonic stem cells. In this context, Lig1 acts as a backup ligase for A-EJ (8) but is redundant for IgH class switch recombination (9, 10). A more recently identified A-EJ component, Polθ, operates on DNA ends in the absence of Ku70/Ku80 (11, 12) and functions independently of Parp1 when repairing G0/G1 DSBs in S-G2/M phase (3). Therefore, it remains unclear which A-EJ mechanisms operate in G0/G1 phase given junction structure and repair capacity differences between noncycling and cycling cells (4).

Microhomology mediated end joining (MMEJ) is not exclusive to a single pathway but rather indicates the extent to which end hybridization was necessary to complete repair (typically 2-20bp). Junctional MHs formed by A-EJ involve limited resection and fill-in processes. They are thought to include, among others, CtIP (13), Mre11 and associated RAD50-NBS1 (MRN) (14–17), and loaders of DNA clamps to tether polymerases, such as Polθ and Polλ (11, 18–22). ATM-mediated phosphorylation of CtIP stimulates sequential Mre11 endo- and exo-nuclease activity to remove protein bound or adducted 5’ ends (23–26) whereas DNA2 promotes resection of clean ends (27). CtIP also promotes DNA2-dependent long-range resection that is separate from an Exo1-dependent mechanism (28–30).

A central regulator of resection that is associated with the full establishment of the DNA damage response (DDR) is 53BP1, which recruits multiple complexes (i.e. Shieldin, CST-Polα, Dynll1 dimers, TOPBP1-ATR, PTIP) to regulate DNA end resection (31–38). Although 53BP1 generally supports NHEJ, it is unclear how these associated complexes support NHEJ. This is further complicated with A-EJ mechanisms which generate resected intermediates to complete repair. Notably, Exo1 is the primary nuclease responsible for long-range resection in G0/G1 phase Lig4-deficient cells, and both Exo1 and DNA2 contribute to long-range resection in cells that are additionally deficient in 53BP1 (39). In this context, long range resection in Lig4/53BP1 double-deficient cells is mediated by ATM (40), which also initiates the DDR with DNA-PKcs, as a functional kinase with Ku70/Ku80, to stabilize end synapsis. While it is clear DSB-associated resection in G0/G1 phase occurs in the absence of core NHEJ factors (4, 39), it is unknown how the DDR regulates A-EJ mechanisms in this context.

Here, we identify the Parp1-XRCC1-Lig3 axis as the primary driver of A-EJ mediated Igκ locus V(D)J recombination in the absence of Ku70. Inhibited V-J recombination due to DDR or candidate repair factor perturbation is accompanied by joints with increased resection and an increased MH over direct joint utilization where ATR, Mre11 exonuclease, polymerase alpha and DNA2 inhibition affect these measures to varying degrees. In this regard, MMEJ utilization becomes near exclusive in *Ku70^-/-^ Xrcc1^-/-^*cells and in *Ku70^-/-^Parp1^-/-^* cells synergized with ATM inhibition. Crucially, we also find A-EJ repair capacity is reliant on 53BP1 in the context of the ATM-initiated DDR. ATM and Parp1 separately support the residual recombination fidelity of coding ends by A-EJ. ATM also suppresses excessive distal V-J recombination and interchromosomal translocations, predominantly to other RAG1/2 DSBs. Other inhibited DDR genes also increase translocations at the cost of V-J recombination efficiency and contrasts that of XRCC1 or 53BP1 deficiencies which are uniformly end joining defective.

## Methods

### Cell Lines

Mouse *vAbl* cells including wild-type (WT, clone B), *L4^-/-^, Ku70^-/-^, and L4^-/-^Ku70^-/-^* (*K7L4^-/-^*) cells were reported in our previous study (4). Putative A-EJ and DDR genes were deleted in the Ku70 deficient (clone B1) or the *Lig4^-/-^Ku70^-/-^*(clone A1-1) *vAbl* lines to generate the following lines: *Ku70^-/-^ Atm^-/-^, Ku70^-/-^Xrcc1^-/-^, Ku70^-/-^53bp1^-/-^, Ku70^-/-^Parp1^-/-^, Ku70^-/-^Exo1^-/-^, Ku70^-/-^53bp1^-/-^Exo1^-/-^* and *Lig4^-/-^ Ku70^-/-^53bp1^-/-^*. Ku70 was ectopically expressed in *Ku70^-/-^*related cell lines using lenti-iKu70-GFP. Confirmation of these cell lines was performed by genotyping using the primers listed in Table S5 and western blotting using the antibody indicated. All *vAbl* cells were cultured at 37°C and 5% CO2 in RPMI-1640 medium supplemented with 10% (vol/vol) FBS, 50 U/mL penicillin/streptomycin, 2 mM L-glutamine, 1× MEM-NEAA, 1 mM sodium pyruvate, 50 μM 2-mercaptoethanol, and 20 mM HEPES (pH 7.4). To induce V(D)J recombination, *vAbl* cells were seeded at a concentration of 1 million cells/ml and supplemented with 3 μM STI-571 for four days. Cells treated with STI-571 were not used for cell line preservation.

### Compounds

The following compounds and their intended effects are indicated: G1/G0 arrest – STI-571 (3μM, TCI Chemicals, Cat # TCI0936-100MG); Cell cycle analysis – EdU (50μM, Cayman Cat # 20518); Inducible gene expression – Doxycycline (2μM, Sigma-Aldrich, Cat # D9891-10G); Parp inhibition – Olaparib (Parpi #1, 10μM (85, 86), AdooQ BioScience, Cat # A10111) and Talazoparib (Parpi #2, 1μM (87,88), ApexBio Cat # A4153); Polθ inhibition – Novobiocin (Polθi #1, 100μM (89), BioVision, Cat # B1526-1G) and ART558 (Polθi #2, 10μM (90), MedChem Express, Cat # HY-141520); Polα inhibition – Adarotene (Polαi #1, 1μM (39, 91), MedChem Express, Cat # HY-14808) and CD437 (Polαi #2, 5μM (92), Sigma-Aldrich, Cat # 178496-5MG); ATR inhibition – AZD6738 (ATRi, 0.5μM (93), MedChem Express, Cat # HY-19323); ATM inhibition – Ku60019 (ATMi, 2μM (94), Sigma-Aldrich, Cat # SML1416-5MG); DNA2 inhibition – C5 (DNA2i, 20μM (95), MedChem Express, Cat # HY-128729); Mre11 inhibition – PFM01 (Mre11eni #1, 10μM), PFM03 (Mre11eni #2, 10μM) and PFM39 (Mre11exi, 100μM) were from the Tainer lab (96).

### Gene Knockouts

The genes indicated in this study were knockout using paired guide RNAs. In brief, 5μl 20μM crRNA were mixed with 5μl 20μM tracrRNA, denatured at 90℃ for 2 mins, and annealed at room temperature for 30 mins to form guide RNA. 5ul 10μM guide RNA mixed with 0.5ul 5xPBS buffer (RNase free) and 0.5ul 60μM SpCas9 protein, incubated at room temperature for 10 mins. Paired guide RNPs were mixed with nucleofection buffer (SF Cell Line 4D X Kit, Lonza, #V4XC-2024) and delivered into 10 million vAbl cells using 4D-nucleofector system (Lonza, Core plus X unit).

### Lentivirus Production

Lentivirus containing ihKu70-GFP-Blast cassette was generated using the 2nd generation lentivirus packaging system as described previously (50). The lentivirus titration was recommended to be performed in the targeting cell lines rather than in HEK293T cells alone. Lentivirus was added into *vAbl* cells and maintained in R10 media with 5μg/ml polybrene for 2 days, changing media and continually cultured in R10 media with 5μg/ml blasticidin for 1-2 weeks (media was changed every 2-3 days to maintain the cell density between 0.1-2 million/ml). The Human Ku70 gene was incorporated into the Lenti-iCas9-neo plasmid (Addgene #85400) through a two-step cloning process. Initially, the neomycin-resistant gene was substituted with a blasticidin-resistant gene. Subsequently, the Cas9 gene was replaced with the Ku70 gene.

### Immunoblotting

The samples, including *vAbl Ku70^-/-^, Ku70^-/-^Xrcc1^-/-^* (#1/2), Ku70^-/-^Atm^-/-^ (#1/2), *Lig4^-/-^ Ku70^-/-^* and *Lig4^-/-^Ku70^-/-^53bp1^-/-^*(#1/2) were collected at a concentration of 5 million cells, spun down and washed by 1ml of RS buffer (150 mM NaCl, 10 mM Tris pH 7.5), spun down and resuspended using 100 μl standard RIPA lysis buffer (150 mM NaCl, 1% NP-40, 0.5% sodium deoxycholate, 0.1% SDS, 50 mM Tris, pH 8.0), leaving on ice for 15mins, then, adding 100 μl of 2X Laemmli buffer (4% SDS, 5% 2-mercaptoethanol, 20% glycerol, 0.004% bromophenol blue, 125 mM Tris pH 6.8) and incubating at 95°C for 10 minutes. Protein electrophoresis, membrane transfer and immunoblotting were performed as previously described (50) using the following antibodies: anti-53BP1 (1:2000, Novus Biologicals #NBP2-54753SS), anti-ATM (1:1000, Proteintech #27156-1-AP), anti-Lig1(1:1000, Proteintech #18051-1-AP), anti-Lig3 (1:5000, BD Biosciences #BD611876), anti-Rabbit-IgG (1:2000, Thermo Scientific #G-21234), and anti-actin (1:4000, Santa Cruz Biotechnology #SC-47778).

### Flow Cytometry Sorting

The *vAbl* cells that transfected with iKu70-GFP was induced by 3μM Doxycycline at least two days before flow cytometry. The population expressed iKu70-GFP was detected in FITC channel and collected in tube with R10 media. The selected cells were cultured for the experiments in this study.

### HTGTS Library Preparation

HTGTS library preparation was performed as previously described (43, 50) with some modifications. In brief, 5.5 μg of genomic DNA from each treatment condition was adjusted to 110μl (50ng/μl) and sheared using a bioruptor sonication device (Diagenode) in low mode for two cycles (30s on + 60s off) at 4°C, resulting in fragments ranging from 200 bp to 2 kb. The sheared fragments were transferred into 96-well microplate and subjected to linear amplification (LAM)-PCR using biotin-labeled primers, including Bio-IgkJ1CE and Bio-IgkJ1SE proximal to the IgkJ1 RAG1/2 incision site, respectively. The LAM-PCR products were enriched using streptavidin-coated 96-well microplate, followed by *in situ* adapter ligation. Unligated adapters were removed, and the ligated products were subjected to nested-PCR using a common primer (AP2I7-novo) matching the adapter sequence and another barcoded I5 primer that matches the region between the bait-site and the biotin-labeled primer. The DNA from the nested-PCR was purified using SPRIselect beads (Beckman #B23318). Subsequently, tagged PCR was performed using primers P7I7 and P5I5, which match the primers used in the nested-PCR. The PCR products were purified by 1% agarose gel electrophoresis, and DNA products with a length of 500 bp to 1 kb were excised and extracted using a Gel-extraction kit. The tagged DNA libraries were subjected to bioanalyzer analysis for quality control and sequencing using the Illumina NovaSeq-PE150. Please refer to Table S6 for the oligos used.

### Data Analyses

LAM-HTGTS data analysis was performed following previously reported methods (4, 43, 50). Briefly, sequencing reads from Illumina NovaSeq PE150 were de-multiplexed based on the inner barcodes and the sequence between the bait site and the nested PCR I5 primers using the fastq-multx tool from ea-utils. The adapter sequences were trimmed using the SeqPrep utility. The demultiplexing and trimming functions were integrated into a script called TranslocPreprocess.pl. Subsequently, the read pairs were normalized down to 500,000 using Seqtk and mapped to the mm9 reference genome using TranslocWrapper.pl to identify chromosome translocations or V-J recombination events, generating result tlx files. Relative translocation measurements were generated like V-J recombination joints by using the no-dedup option from TranslocWrapper.pl (43, 97), whereas absolute translocation measures for junction structures were derived using default settings (4, 43). Junctions that aligned to the bait region were not shown in the result tlx files and were extracted separately using a script called JoinT.R, resulting in JoinT tlx files containing translocations, V(D)J recombination and rejoin events of IgkJ1 region.

JoinT tlx files were converted into bedgraph files using tlx2bed.py, which were then visualized and plotted using IGV (integrative genomics viewer). Junctions in regions of interest from the JoinT tlx files were extracted using tlxbedintersect.py, which relied on two other scripts, tlx2BED.pl and pullTLXFromBED.pl. The regions of interest varied depending on the specific questions. For Vκ-Jκ recombination, the regions of interest were the RAG1/2 cleavage sites of Vκ genes, with a flanking 200 bp window (± 200), that could be further divided into 4-5 topologically associating domains (TADs). In the case of translocation, the region of interest was the “prey” locus in all chromosomes except chr6. To visualize the relative translocation distributions, the representative JoinT tlx files were converted into circos plots.

The JoinT tlx files were also utilized for repair efficiency, pathway, and resection analyses. V-J efficiency (Vs% in Tables S3, S4) was calculated by dividing the junctions recovered from the Vκ region by the normalized total reads. JctStructure.R was employed to determine the repair patterns, including microhomology (MH), direct repair, and insertion over a range of 20bp for each extreme. The degree of DSB end resection, indicated by the distribution of junctions near the DSB break site, was quantified using ResectionRSS.R.

Data obtained from western blot and flow cytometry experiments were analyzed using ImageJ (NIH) and FlowJo (FlowJo, LLC), respectively.

### Software/Code Availability

All essential HTGTS-specific codes are described elsewhere (4, 43, 50) or are publicly available at https://github.com/JinglongSoM/LAM-HTGTS. The codes and software used in this study are indicated: Python (v3.8.5); R (v4.0.3); ImageJ; IGV (v2.8.2); FlowJo (v10.8)

### Statistical Analysis

Data are reported as mean ± SEM unless specified otherwise. Differences were analyzed using one-way and two-way ANOVA, followed by Dunnett’s multiple comparisons test. Statistical calculations were carried out using GraphPad Prism 10 (GraphPad Software, Inc.). A P-value of less than 0.05 was deemed statistically significant.

## Results

To elucidate mechanistic factors for Ku-independent A-EJ and their regulation, we knocked out several DDR or candidate A-EJ genes in the *Ku70^-/-^*, *Ku70^-/-^53bp1^-/-^*, and *Lig4^-/-^Ku70^-/-^* Abelson kinase transformed murine progenitor B (*vAbl*) cell lines and validated low levels of prior endogenous V(D)J recombination (Fig. S1A-J; Tables S1-S4). Ku70 was complemented to restore NHEJ and contrast A-EJ phenotypes. We used a panel of common inhibitors of implicated kinases, polymerases, and nucleases in both WT and *Ku70^-/-^* backgrounds to determine the core A-EJ pathway and its regulators (see methods for cited doses). *vAbl* cells were assayed for physiologic V(D)J recombination under G0/G1 arrest by the Abl kinase inhibitor STI-571 (4, 40, 41, 42) and for cell viability to optimize compound efficacy and interpretation of recombined junctions. Overall, most inhibitors exhibited minimal effects, though Polα inhibition had a discernable impact on cell viability (Fig. S2A-L).

We measured Igκ locus V-J recombination and interchromosomal translocations by A-EJ in *Ku70^-/-^*and derivative *vAbl* cells using the high throughput rejoin and genome-wide translocation sequencing platform, *HTGTS-JoinT-seq* (4, 43), from the Jκ1 bait DSB position (Fig. S3A). We employed baits from each side of the DSB corresponding to both the hairpin sealed coding end (Jκ1CE) and blunt signal end (Jκ1SE), a result of paired RAG1/2 DSB cleavage in complex with one of 100+ corresponding Vκ coding/signal end preys. This locus also contains Vκ gene segments organized in deletional (DEL) or inversional (INV) strand orientations with respect to the Jκ region that can result in a variety of deletions, inversions and excision circles when viewed from the coding or signal ends of the Jκ1 bait DSB (Fig. 1A, S3A, S4A). In general, although chemical or genetic perturbation of A-EJ yielded similar results for both bait ends, with some exceptions highlighted below, signal ends tended to recover more junctions than coding end baits. The same outcome did not occur for NHEJ, indicating that hairpin opening, an end processing event that is required for accessibility and subsequent ligation of coding ends, may be rate limiting.

**Figure 1.**
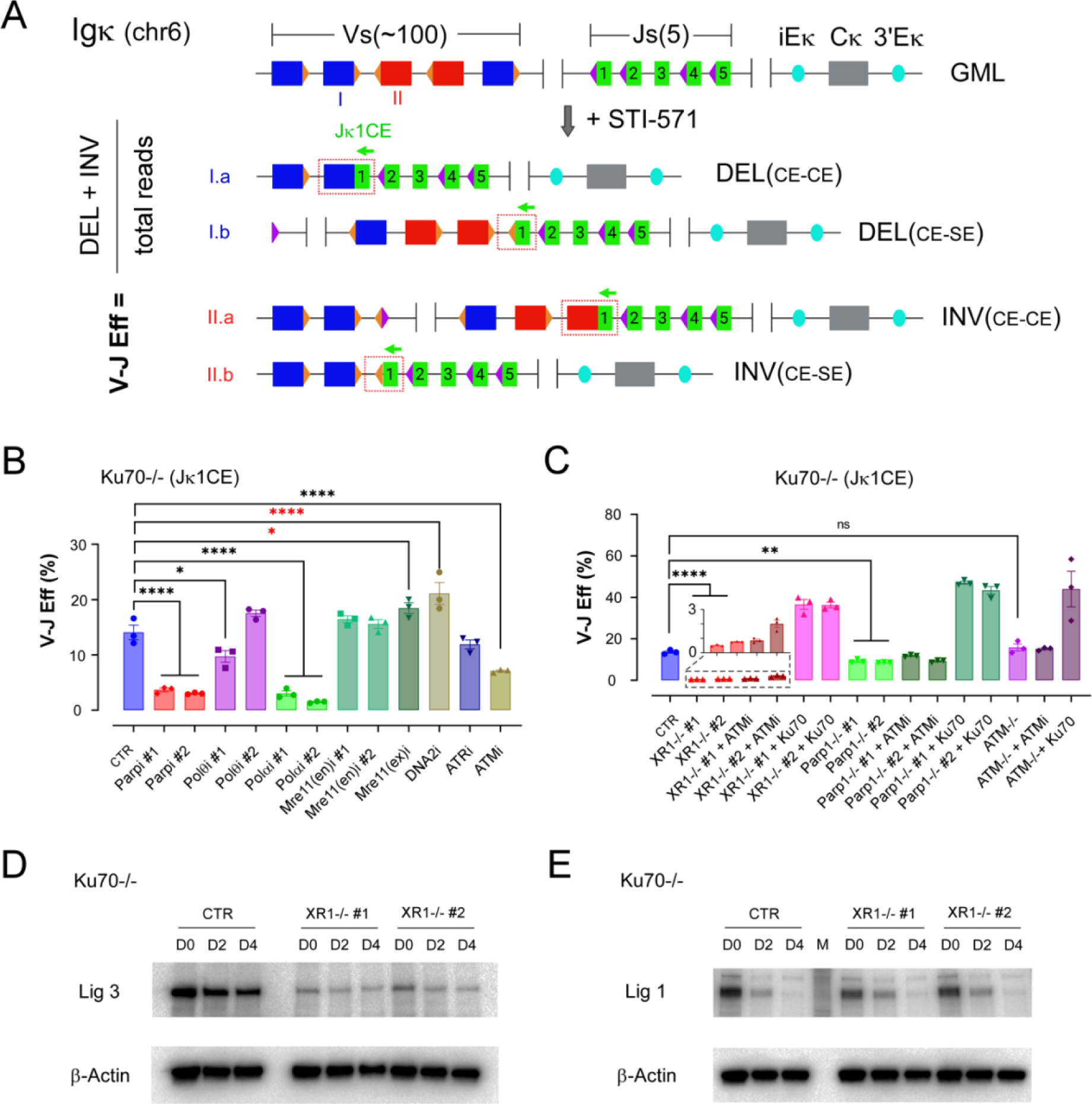
XRCC1, Parp1, and DDR components support A-EJ of Igκ locus DSBs. (**A**) The murine Igκ antigen receptor locus in a germline configuration (GML), each with an associated recombination signal sequence (triangles). STI-571 treatment enables *vAbl* cells to undergo G0/G1 arrest and initiate V-J recombination. V gene segments are oriented in a deletion (DEL, blue; example: I) or inversion (INV, red; example: II) configuration with respect to the Jκ1 coding end (CE) bait (green arrow). The Jκ1CE can form Vκ coding (CE-CE; I.a or II.a) or hybrid (CE-SE; I.b or II.b) joints with the associated recombination signal end (SE; orange triangles). Recombination efficiency (V-J Eff) was calculated by the sum of DEL and INV V-J joints divided by the total reads. (**B**) V-J recombination efficiency of *Ku70^-/-^*cells with or without inhibitors. (**C**) V-J recombination efficiency changes of *Ku70^-/-^ vAbl* cells with added deletions and optionally with ATMi treatment or Ku70 ectopic expression. All experiments were biologically repeated three times, and significance was determined by one-way ANOVA with post-test comparisons: * (p<0.05), ** (p<0.01), *** (p<0.001), **** (p<0.0001) and ns (no significance); red asterisks indicate significant increases. (**D**) Western-blot of Lig3 expression levels in *vAbl Ku70^-/-^* (CTR) and *Ku70^-/-^ XRCC1^-/-^*(*XR1^-/-^* #1/2) cells that treated by STI-571 for 0, 2 and 4 days (D0, D2, D4), respectively, where β-Actin used as controls. (**E**) Same as (D) but for Lig1 expression levels.

### XRCC1, Parp1, and DDR components support A-EJ of Igκ locus DSBs

As A-EJ recombines Vκ-Jκ regions with significantly less bias (i.e. CE-CE and SE-SE) than NHEJ (4), we incorporated repair to both CEs and SEs in the Vκ region, normalized against the total sequence reads, which include other repair outcomes (e.g. rejoined or recombined Jκ DSBs and translocations), to derive the V-J recombination efficiency (V-J Eff)(Fig. 1A, S4A)(see methods). Using this approach, we separated analysis of gene perturbations in the *Ku70^-/-^* parental background into two distinct groups: putative core factors (Parp1, Polθ, and XRCC1) and DDR factors involved in regulation (ATM and ATR) or DNA end processing (Polα, Mre11, and DNA2).

Regarding putative core factors, we found inhibition of multiple Parp genes (Parpi #1/2) or deletion of Parp1 (*Parp1^-/-^* #1/2) decreased V-J efficiency by 80% and 30% for the CE bait and 85% and 80% for the SE bait, respectively (Figs. 1A-C, S3B, S4B-D; Tables S3, S4), indicating Parp genes play a significant role in this process. We next wanted to know whether Polθ participates in this mechanism despite its very poor expression in G0/G1 phase (3). Here, Polθi #2 had no impact on V-J recombination efficiency (Figs. 1B, S4C); however, Polθi #1 decreased both WT and *Ku70^-/-^* recombination (Figs. 1B, S4C, S5B-C). Given that Polθi #1 (Novobiocin) also affects ATPase activities of multiple classes of proteins (44), some of which may be important for recombination, whereas Polθi #2 (ART558) is, to date, more extensively characterized (45), we propose that Polθ plays a minimal role at best in Ku-independent V(D)J recombination. We next determined which A-EJ components drive ligation. As nuclear Lig3 activity depends on XRCC1 (9, 46), and likewise for Lig1 with PCNA (47), we compared total protein levels of the A-EJ ligases in cycling versus G0/G1 arrest. Lig3 abundance was only slightly decreased (Fig. 1D), whereas Lig1 levels plummeted 5-fold by day 2, when cells are arrested, and was nearly absent by day 4 (Fig. 1E). This suggests Lig3 is the primary A-EJ ligase in this G0/G1 setting. As predicted from cycling cells (48), XRCC1 deletion reduced steady-state protein levels of Lig3 by 6-fold but did not affect Lig1 (Fig. 1D-E), indicating XRCC1 specifically stabilizes Lig3 protein levels. In this regard, XRCC1 deletion essentially abolished V-J recombination resulting in a robust 1-2 order of magnitude decrease (Figs. 1C, S3B, S4B, S4D), revealing a core A-EJ role for XRCC1/Lig3. Collectively, the data support a Parp1-XRCC1-Lig3 axis driving G0/G1 phase A-EJ.

With regard to the regulation of A-EJ, both ATR and ATM inhibition displayed modest reductions (∼40-50%) in V-J recombination (Figs. 1B-C, S3B, S4B-C), indicating DDR kinase activities partially facilitate A-EJ. Intriguingly, although ATMi treatment of *Ku70^-/-^Atm^-/-^ vAbl* cells had no additional impact to V-J efficiency (Figs. 1C, S4C), deletion of ATM in *Ku70^-/-^* cells did not decrease V-J efficiency as it did with ATMi alone. This suggests a dominant negative activity suppresses overlapping DDR kinase functions. As expected, NHEJ restoration via Ku70 expression in double knockout cell lines dramatically increased V-J efficiency, reaching levels 2-fold higher than *Ku70^-/-^* alone (Figs. 1C, S4C). Inhibiting the nuclease activity of DNA2 displayed opposing efficiency changes that was dependent on the bait end. Specifically, DNA2i decreased efficiency, on par with ATRi for the SE bait, but increased efficiency by ∼33% for the CE bait, which was the greatest increase of all combinations tested. This increase with DNA2i suggests competition with another factor that opens hairpins. Inhibited Mre11 nuclease activities had a smaller impact on recombination efficiency, where inhibited exonuclease activity (Mre11(ex)i) modestly increased CE bait V-J efficiency, like DNA2i, but did not affect the SE bait, unlike DNA2i (Fig. 1B, S4C). Mre11 endonuclease inhibitors (Mre11(en)i #1/2) had no significant impact on any analysis parameter in this study. These results are in support of a prior *in vitro* finding that DNA2, rather than Mre11, acts as the primary nuclease for accessible ends (27).

To uncover additional insight into the chemical inhibitory effects specific to A-EJ, we also tested the same compounds in *WT vAbl* cells. Parpi #1/2, ATRi and ATMi compounds also reduced V-J efficiency from both CE and SE baits (Fig. S5A-C), but the severity was attenuated compared to *Ku70^-/-^* cells (Fig. 1B, Fig. S4C). Polα inhibition (Polαi #1/2) displayed a 2-3 fold reduced V-J efficiency in both WT and *Ku70^-/-^ vAbl* lines; however, the decreased viability (Fig. S2A-C, S2G-I) limits interpretation of junction yields, but distinguishing repair patterns are evident and described below. Overall, the data suggest an ATM/ATR DDR mechanism supports A-EJ.

### 53BP1 is essential for G0/G1 A-EJ

We next addressed the extent to which the DDR supports A-EJ by deleting a core regulator of resection: 53BP1. Loss of 53BP1 (*Ku70^-/-^53bp1^-/-^* #1/2) inhibited A-EJ, resulting in a robust decrease (10-fold for CE bait; 5-fold for SE bait) in V-J efficiency for all combinations (Figs. 2A-B, S6A-B, S7A-C; Tables S3, S4) but was still comparably higher than *Ku70^-/-^Xrcc1^-/-^* (#1/2) cells across experiments. As Exo1 is a crucial nuclease responsible for long-range resection in *vAbl* cells without 53BP1 (39), we generated *Ku70^-/-^53bp1^-/-^Exo1^-/-^* (#1/2/3) cells to determine if recombination efficiency was restored. Unexpectedly, no dramatic improvement was detected (Fig. S6A-B), suggesting something beyond hyper resection control is necessary to restore V-J efficiency. Similarly, Exo1 deletion in *Ku70^-/-^* cells did not alter the recombination efficiency (Fig. S8A-B), and DNA2i or Mre11(ex)i treatments to *Ku70^-/-^53bp1^-/-^*(#1/2) and *Ku70^-/-^53bp1^-/-^Exo1^-/-^* cells (#1/2) also did not significantly change recombination efficiencies (Fig. S6B). Therefore, we conclude Exo1 deficiency alone or in combination with inhibited DNA2 or Mre11 exonuclease activities cannot restore A-EJ-mediated V-J recombination efficiency in the absence of 53BP1.

**Figure 2.**
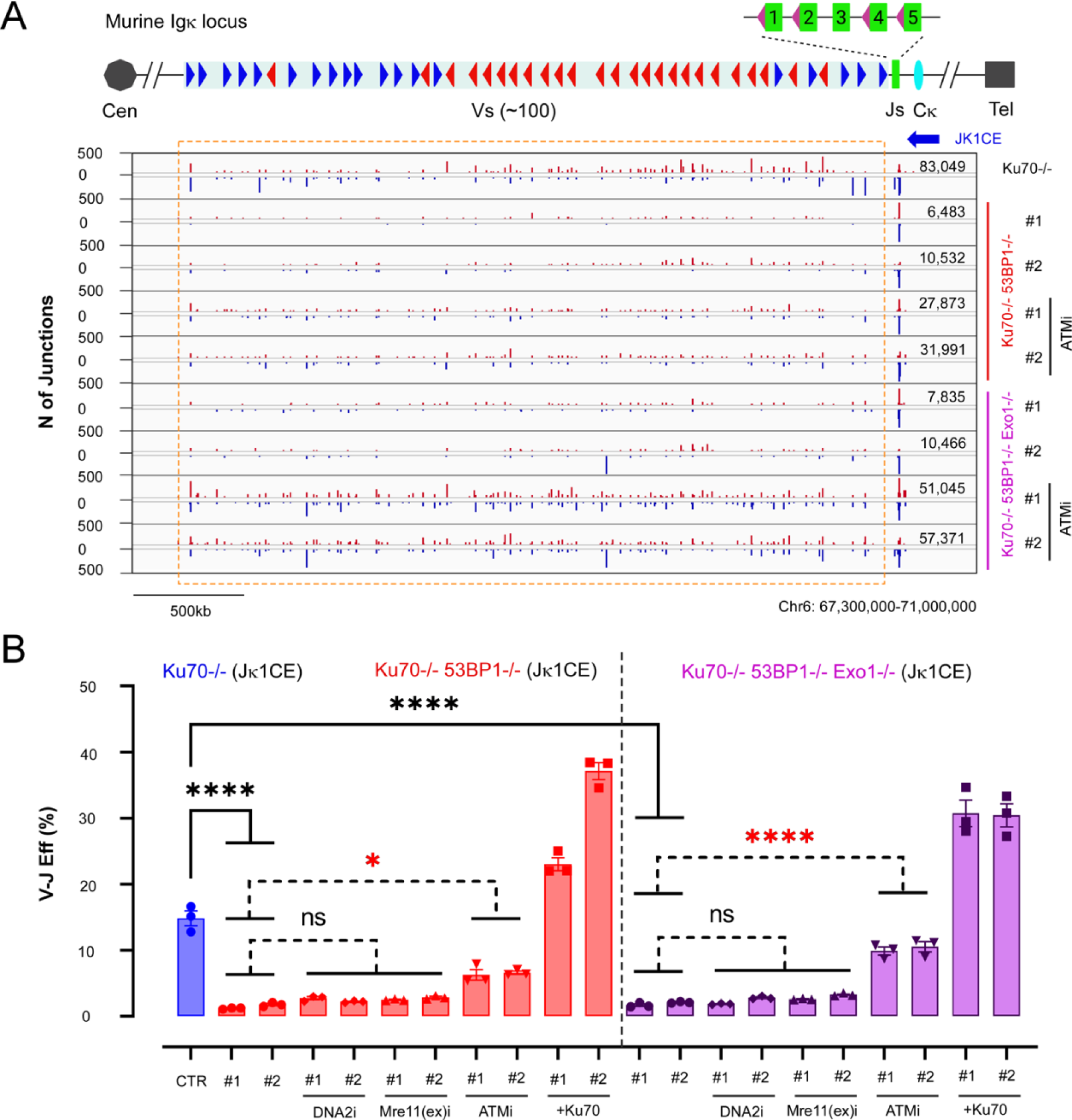
53BP1 is essential for G0/G1 A-EJ. (**A**) Representative *Ku70^-/-^* CE bait junction plots as described in Fig. 1 but in the context of 53BP1 single or 53BP1/Exo1 double deletion, with or without ATM inhibition. (**B**) V-J recombination of the above backgrounds, with or without DNA2, Mre11(ex) and ATM inhibitors, or Ku70 rescue expression. Differences within *Ku70^-/-^53BP1^-/-^* (red bars) and *Ku70^-/-^53BP1^-/-^Exo1^-/-^*(magenta bars) are evaluated by two-way ANOVA plus post-comparison: * (p<0.05), **** (p<0.0001) and ns (no significance); black asterisks indicate significant decreases. All experiments were biologically repeated three times.

ATM inhibition in G0/G1-arrested *Lig4^-/-^53bp1^-/-^ vAbl* cells blocks long-range resection (49). Therefore, we tested ATMi effects in *Ku70^-/-^53bp1^-/-^* (#1/2) and *Ku70^-/-^53bp1^-/-^Exo1^-/-^* (#1/2) cells and found V-J efficiency from CE and SE baits was restored to ∼50% and ∼80% of that in *Ku70^-/-^*, respectively, but comparable to ATMi-treated *Ku70^-/-^* (Fig. 2A-B, S6A-B). Correspondingly, ATMi decreased V-J efficiency in *Ku70^-/-^Exo1^-/-^*and *Lig4^-/-^Ku70^-/-^* cells (Figs. S7B-C, S8A-B). Together, the data indicate ATM inhibition, which partially disrupts A-EJ, can normalize the impact of 53BP1 deficiency on A-EJ. Restoring DNA-PK functionality via Ku70 complementation irrespective of 53BP1 status enhanced recombination efficiency beyond *Ku70^-/-^* except in Lig4 deficient backgrounds which remained unchanged (4, 50) (Figs. 2B, S6B, S7B-C, S8A-B; Tables S3, S4). In sum, we conclude 53BP1 robustly supports A-EJ-mediated recombination in the context of the ATM-activated DDR.

### A-EJ regulators and drivers, but not 53BP1, suppress resected end joining

An increased range of resected joints relative to NHEJ-mediated joints (4) serves as a distinguishing feature of A-EJ, which is kinetically relevant to their inherent repair efficiencies. Thus, we wanted to know which of the above perturbations affected resected joint distributions. To do this, Vκ joints were pooled and remapped to an absolute DSB position located between the prey coding end and its adjacent signal end for both Jκ1CE and Jκ1SE baits. In *WT vAbl* cells, SE baits were joined to SE preys, likewise for CE baits/preys, and were exclusive to a ± 10bp window around the Vκ DSB sites (Fig. S9A-C). In contrast, *Ku70^-/-^* CE and SE baits each contained a mix of CE and SE prey joints with distributions that extended far beyond this window (Figs. 3A, S10A-D). Therefore, we derived the fraction enriched within the ± 10bp DSB window to discern changes in resected joint distributions. Despite varied effects on V-J efficiencies with the inhibitor panel in WT cells (Fig. S5B-C), none of them promoted resected end joining (Fig. S9B-C). In contrast, Parp (#1/2), Polα (#1/2), and ATM inhibitors increased the fraction of resected joints in *Ku70^-/-^* cells (Fig. 3A-B, S10A-B). *Parp1*^-/-^ (#1/2), *Atm^-/-^*, or *Xrcc1^-/-^* (#1/2) in *Ku70^-/-^*cells also significantly increased resected joints (Fig. 3C, S10C). However, corresponding 53BP1, Lig4, and/or Exo1 deletions in *Ku70^-/-^* cells did not increase the resected joint fraction unless they were additionally treated with ATMi (Figs. 3D, S10D, S11A-F). In contrast, ATMi did not significantly change joint distributions when added to *Ku70^-/-^Parp1^-/-^* (#1/2) or *Ku70^-/-^Xrcc1^-/-^* (#1/2) *vAbl* cells (Figs. 3C, S10C). This suggests Parp1 and XRCC1 have overlapping roles with ATM to regulate end processing. Neither DNA2 or Mre11 nuclease inhibition nor Exo1 deletion displayed an anti-resected joint enrichment in the various backgrounds (Figs. 3A-B, 3D, S9B-C, S10B, S10D, S11C-F), except for DNA2i in *Ku70^-/-^* cells (Figs. 3A-B, S10A-B), which correlates with its impacts on V-J recombination efficiency. As predicted, restoration of NHEJ through Ku70 expression dropped the resected joint fraction by ∼2-fold in relevant cell lines except those with additional Lig4 deficiency (Figs. 3C-D, S10C-D, S11A-F; Tables S3, S4), which are subject to DNA-PK-mediated resection mechanisms (50). Collectively, we conclude that DNA2 promotes resected end joining while Parp1, XRCC1, Polα, and ATM signaling limit resected end joining in the absence of Ku70. We also conclude, counterintuitively, that 53BP1 does not influence resected joint distributions despite crucially supporting A-EJ recombination efficiency.

**Fig. 3.**
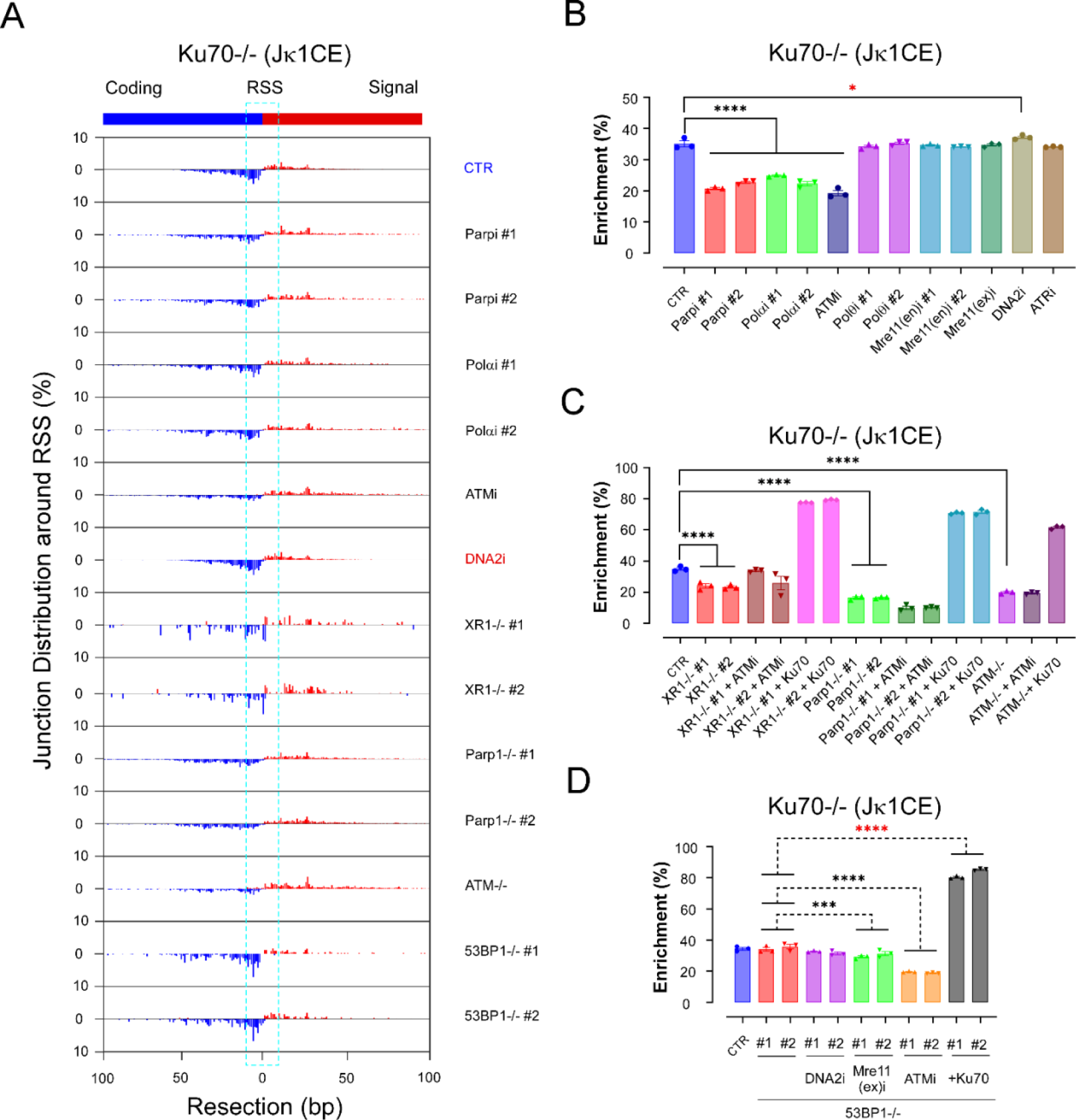
A-EJ regulators and drivers, but not 53BP1, suppress hyper-resected joints. (**A**) Coding and signal prey junctions of Vκ region DSBs from the Jκ1CE bait are aggregated in a resection window of ±100 bps around the RAG1/2 DSB (RSS). Representative plots showing restricted (DNA2i treatment), extended (Polα inhibition; XR1 deletion; Parp1 and ATM inhibition or deletion), or no significant change (53BP1 deletion) are indicated. (**B-D**) The percentage of junctions that enriched in the window of ±10 bps around RSS site was used as indicator for resection, as cyan dashed rectangle in (A). The significance of enrichment changes when combined with indicated inhibitors or gene modification or both were evaluated by one-way ANOVA plus post comparison (B-C) and two-way ANOVA plus post comparison (D); N=3.

### MMEJ increases with greater declines in A-EJ activity

As resection enhances the likelihood of relying on junctional MHs to stabilize/align ends and complete ligation, we wanted to know whether increased resected joints correlated with changes in Vκ region junction structure patterns. For a baseline A-EJ measure, *Ku70^-/-^* joints from either CE or SE baits contain very few insertions, possess a dominant peak of direct joints (∼50-60%), along with an exponential decay of increasing MH lengths (1-4 bps)(Fig. 4A). In contrast, NHEJ restoration via Ku70 complementation reduced MH use and increased direct repair in general and specifically increased palindromic insertions from CE baits, consistent with what is observed in WT *vAbl* cells (4, 50, 51) (Fig. 4A-D, S12A-F, S13A-B, S14A-D, S15A-D). For the A-EJ perturbations, both CE and SE bait junctions displayed similar pattern changes across comparisons with the 1bp MH representing an inflection point from the *Ku70^-/-^* joint structure pattern. Of the 12 inhibitors, only ATM and Polα (#1/2) inhibitors increased total MMEJ (>1bp) by ∼100% while Parp (#1/2) inhibitors marginally increased MMEJ. *Ku70^-/-^Atm^-/-^* cells also reproduced the phenotype of ATMi-mediated MMEJ increase. Although *Ku70^-/-^Parp1^-/-^* (#1/2) structures were not altered, inclusion of ATMi promoted MMEJ synergy with a 30-40% transition of direct to MH joint utilization (Figs. 4A-B, S12A-D). However, when integrated with an unchanged resected joint pattern with combined Parp1 and ATM perturbation (Fig. 3C, S10C), the data suggests resection alone is not causal for the MMEJ synergy. Furthering this notion, *Ku70^-/-^ 53bp1^-/-^* (#1/2) lines, which did not increase the range of resected joints, increased total MMEJ to a similar level as ATMi alone. Added Mre11, DNA2 or ATM inhibitors did not affect the increased MMEJ pattern (Figs. 4C, S12E). A similar result for these inhibitors was observed in other 53BP1 deficient backgrounds, and Exo1 deficiency in the various backgrounds did not enhance MMEJ (Fig. S13A-B, S15A-D). A robust transition to MMEJ that was similar in phenotype to *Ku70^-/-^Parp1^-/-^* + ATMi was revealed in *Ku70^-/-^Xrcc1^-/-^* cells (#1/2) which used ∼18% direct joints. In this regard, direct joint utilization was further reduced to ∼8% with ATMi and is reminiscent of the G0/G1 phase *Lig4^-/-^ vAbl* MMEJ pattern for both bait ends (50). Collectively, these observations imply increased MMEJ does not necessarily rely on altered resected joint patterns, but rather, the altered patterns may be a consequence of increased MMEJ utilization. The data also supports the concept that near-exclusive MMEJ utilization is suppressed by both the ATM-recruited 53BP1 chromatin domain and the *bona fide* A-EJ pathway.

**Fig. 4.**
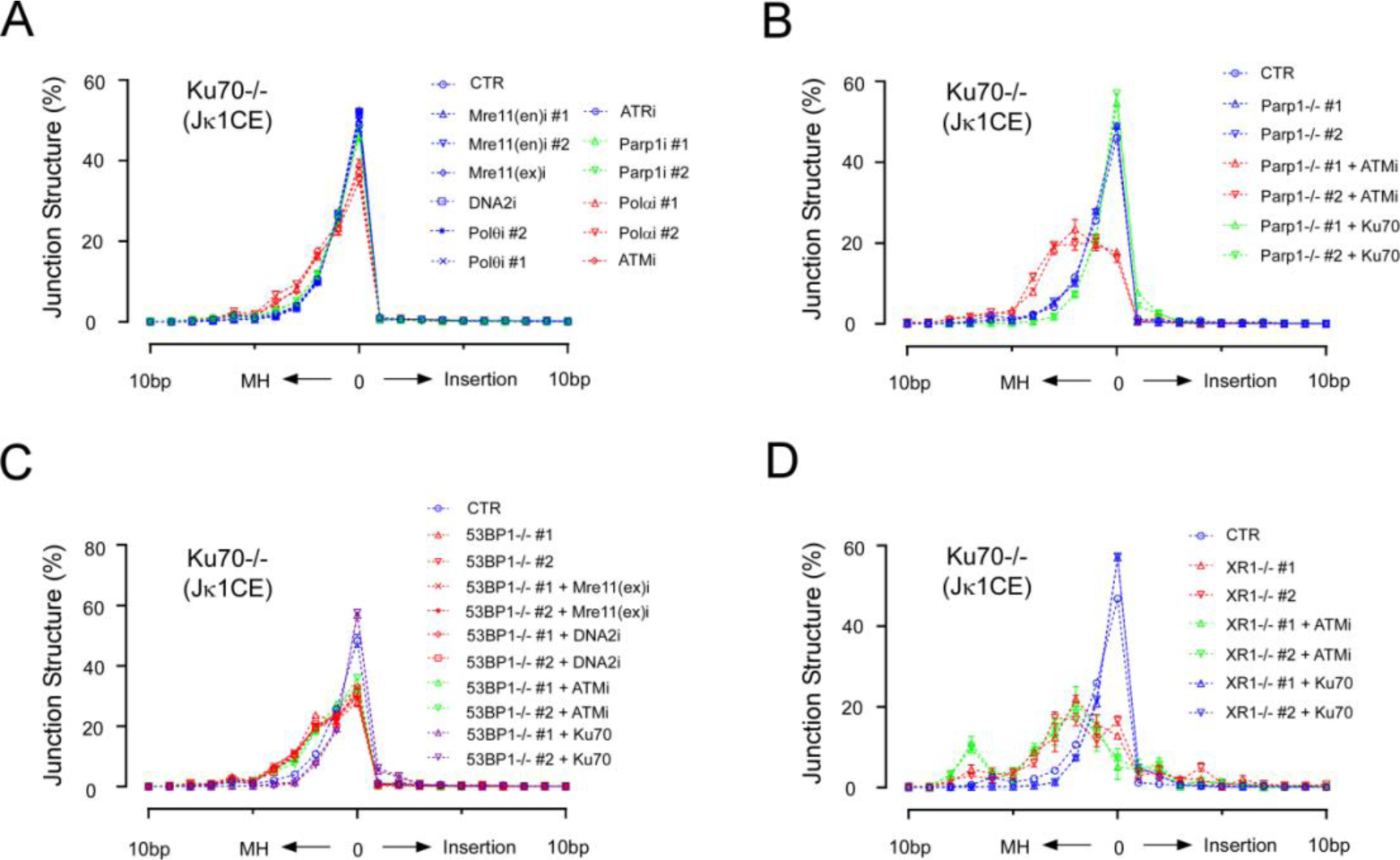
MMEJ increases with greater declines in A-EJ function. (**A**) Jκ1CE bait and Vκ region prey junction structure distributions for the various inhibitors are split into three groups: no detectable pattern change (blue), marginal change (Parpi #1/2, green) and significant change (Polαi #1/2 and ATMi). The repair profile was shown in ±10bp window, the left part indicates overlapping bait/prey microhomology (MH) length, 0 indicates direct (blunt) repair and the right part indicates the insertion size. (**B-D**) Same as (A) but for Parp1, 53BP1 and XRCC1 deletion with/without indicated inhibitors or Ku70 expression, respectively. All experiments were biologically repeated three times.

### ATM and Parp1 maintain recombination fidelity

Given that the DDR supports A-EJ functions, we evaluated whether perturbing pathway components altered Vκ region CE and SE pairing to the Jκ1 bait DSB to further promote CE-SE hybrid joining. To do this, we quantified the number of junctions based on the joining types and derived the CE/SE ratio (Fig. 5A; Tables S3, S4). From the CE bait in WT *vAbl* cells, CE-CE recombination fidelity averaged ∼850:1, and ATMi decreased bias ∼18-fold, down to ∼45:1 (Fig. S16A; Table S3) but remained dependent on NHEJ (52). Among the other inhibitors, only Parpi (#1/2) displayed an increase in these hybrid CE-SE joints, but this effect was only ∼2-fold. Although the absence of Ku70 significantly drops CE recombination fidelity down to ∼2:1 (4, 50), we discovered that inhibited ATM and Parp had similar effects in further decreasing recombination fidelity toward a true translocation-defined 1:1 CE/SE ratio with no additional inhibitors impacting recombination fidelity (Fig. 5B). Similar to the trend of increased resected joints (Figs. 3A-D), *Parp1^-/-^* (#1/2), and *Xrcc1^-/-^*(#1/2), but not *53bp1^-/-^* (#1/2) or *Exo1^-/-^* (#1/2/3) also decreased the residual end bias (Figs. 5B-C, S16B-C). However, while *Ku70^-/-^Atm^-/-^*or *Ku70^-/-^* + ATMi dropped CE bias down to ∼1:1, SE bias from the Jκ1SE bait increased ∼3-fold to ∼3:1 (Fig. 5B-C; Tables S3, S4); this ATMi effect to recover SE fidelity was also found across all combinations, including 53BP1 deficiency (Table S4). This is notable since only ATM perturbation displayed both a CE bias drop and a SE bias rise in A-EJ backgrounds (Tables S3, S4), supporting prior work that ATM regulates RAG1/2 post-cleavage activity in DNA-PK deficient cells (53).

**Fig. 5.**
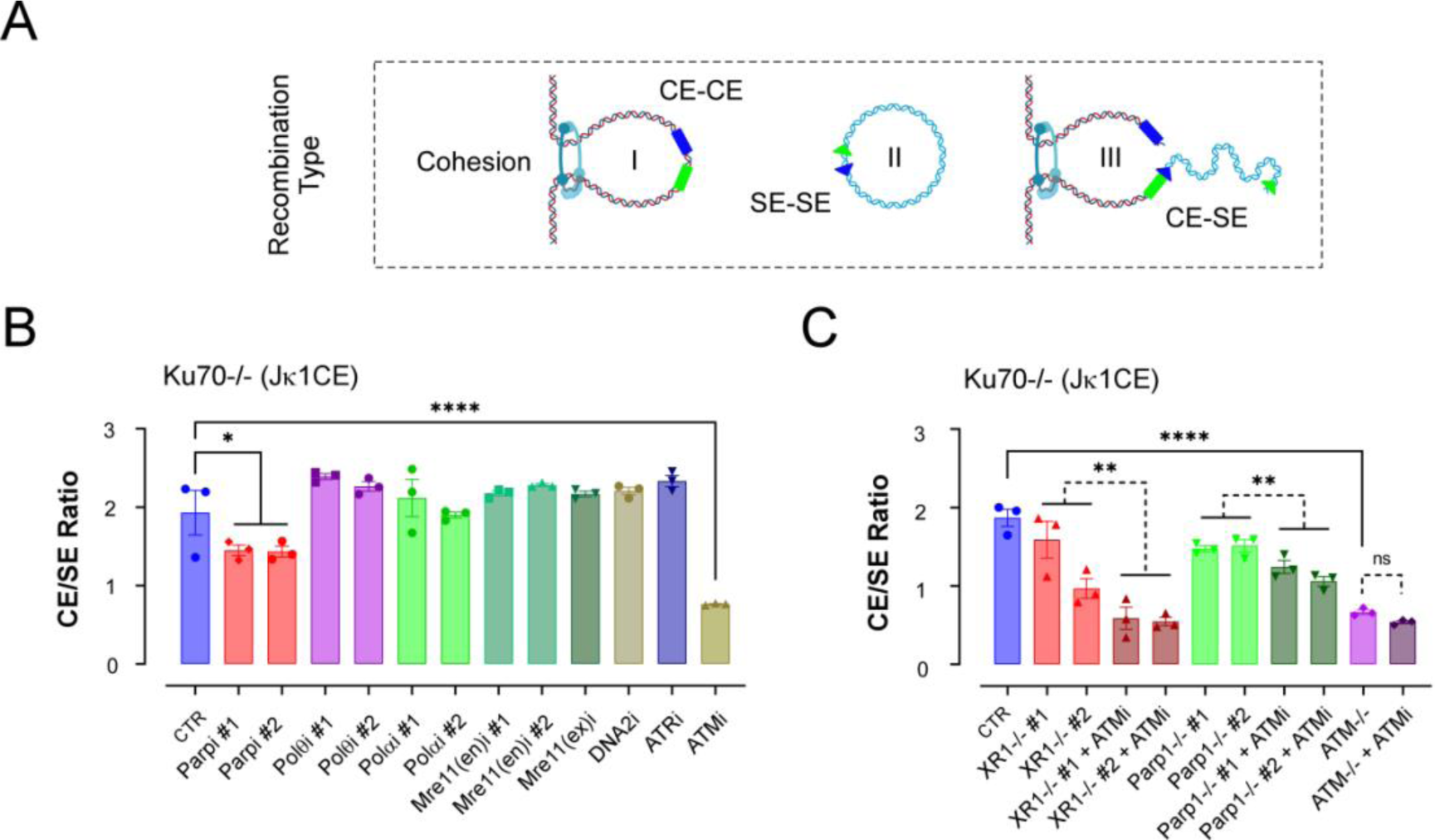
ATM and Parp1 maintain recombination fidelity. (**A**) An illustration of possible recombination outcomes of post-RAG1/2 cleavage, including coding to coding (CE-CE) (I), signal to signal (SE-SE)(II), and hybrid joins (CE-SE)(III). (**B**) The *Ku70^-/-^* cell CE/SE ratio from Jκ1CE bait are indicated with/without indicated inhibitors. (**C**) Same as (B) but with additional XRCC1 (XR1), Parp1 or ATM deletion with/without ATM inhibitor. One-way ANOVA with post-test significance for each comparison are indicated; N=3.

### A-EJ and ATM increase distal Vκ recombination

The Igκ topologically associating domain (TAD) can be divided into Vκ region sub-TADs – sTAD1-2, sTAD3, sTAD4, and sTAD5 (54), each with varying numbers of actively extruding CTCF/cohesion anchored chromatin loops which form an interaction zone for V-J pairing and cleavage by RAG1/2 (55). In *WT vAbl* cells, Jκ1CE bait recombination was the greatest in the Jκ proximal sTAD (sTAD5; ∼45%) and declined as a function of linear distance with the distal sTAD (sTAD1-2) contributing ∼11% of total recombination (Fig. S17A-B; Table S3). In contrast, *Ku70^-/-^ vAbl* cells displayed a similar overall trend but with ∼7% decrease in sTAD5 and ∼7% increase in sTAD1-2 (Fig. 6A-C; Table S3). From the panel of inhibitors tested, none displayed a significant shift in the landscape of utilization from WT cells (Fig. S17B; Tables S3, S4). However, only ATMi or *Atm^-/-^*further increased sTAD1-2 recombination in *Ku70^-/-^* by ∼5-6% (Figs. 6B-C, S18A-C; Table S3) and was consistently found across other *Ku70^-/-^* combinations except for *Ku70^-/-^Xrcc1^-/-^* (Figs. 6C, Tables S3, S4). We conclude A-EJ and inhibited ATM have additive effects on promoting distal V-J recombination.

**Fig. 6.**
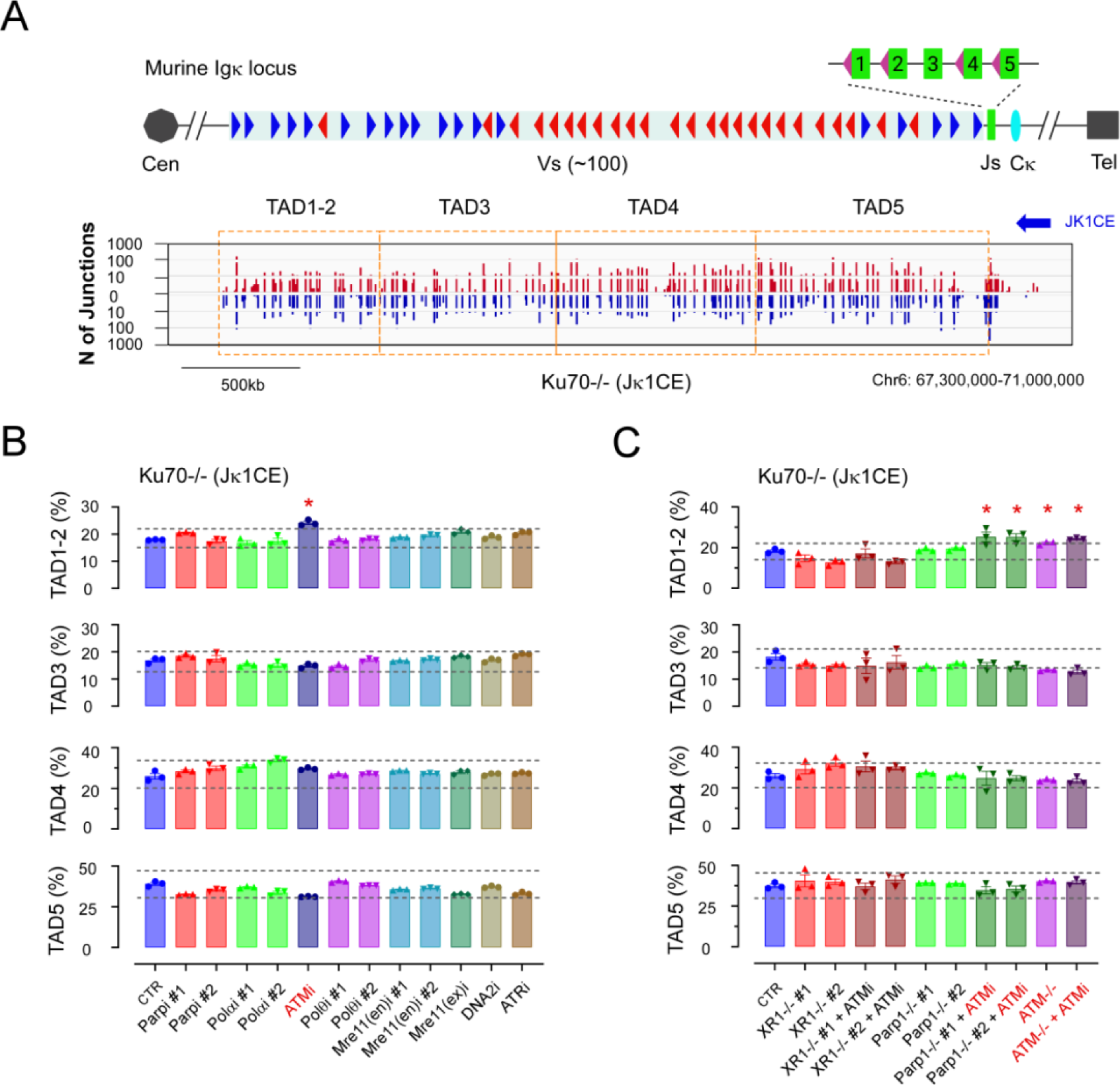
A-EJ and ATM increase distal Vκ recombination. (**A**) The Igκ locus is a topologically associated domain (TAD) that can be divided into four subTADs: sTAD1-2, sTAD3, sTAD4 and sTAD5 (54), as shown in the *Ku70^-/-^* Jκ1CE bait control. (**B**) The percentage of the four segments with or without indicated inhibitors as indicated. (**C**) Same as in (B) but for XRCC1 (XR1), Parp1 and ATM deletions with or without ATM inhibitor. Text highlighted in red, and indicated in the graph by red asterisk are conditions that extend beyond the ± 20% threshold change of the CTR (dashed lines). All experiments represent three biological replications.

### Compromised DDR affects A-EJ translocations

Despite the normally well-orchestrated process of V(D)J recombination, chromosome translocations between RAG1/2-mediated and spontaneous DSBs can also occur. Thus, we sought to determine whether genome-wide junctions are repaired similarly to recombined junctions in the Igκ locus in the context of A-EJ and DDR gene perturbation. In this regard, translocations represented a mix of spontaneous prey DSBs and RAG1/2 prey DSBs from other antigen receptor loci (e.g. Igλ, IgH, etc.). Therefore, we analyzed both the relative and absolute translocation pools (see methods), which were highly correlated (Tables S3, S4), to describe significant changes. In general, WT translocations were low in frequency and composed mostly of spontaneous DSB end partners. Whereas *Ku70^-/-^* translocations were 3-to 6-fold higher in frequency with a greater proportion coming from other antigen receptor locus RAG1/2 DSB end partners (Figs. 7A, S19A, S20A; Tables S3, S4). Consistent with the negative impact to V-J efficiency (Fig. S4B-C), WT *vAbl* cells treated with Parp, ATM, or ATR inhibitors increased translocations (Fig. S19A-H). Notably, *Ku70^-/-^ vAbl* cells treated with Parpi (#1/2) specifically increased spontaneous translocations, while ATMi preferentially increased RAG1/2 translocations (Fig. S19B-D; Tables S3, S4). This difference emphasizes their respective roles in base excision repair and regulating RAG1/2 recombination center activity. Unlike WT CE bait V-J joints which were predominantly direct with some insertions (∼20%), WT CE bait translocation joints were primarily insertions (∼70%) with a cascading enrichment of direct (∼20%) and MHs (∼10%). This pattern is consistent with coding end preservation and committed repair by NHEJ. In this context, many of the inhibitors increased direct and MH utilization, where the compounds that decreased V-J efficiency and increased translocations, like ATMi, Parpi, ATRi, and Polαi, also displayed the greatest MMEJ transition (Fig. S19I-J).

**Fig. 7.**
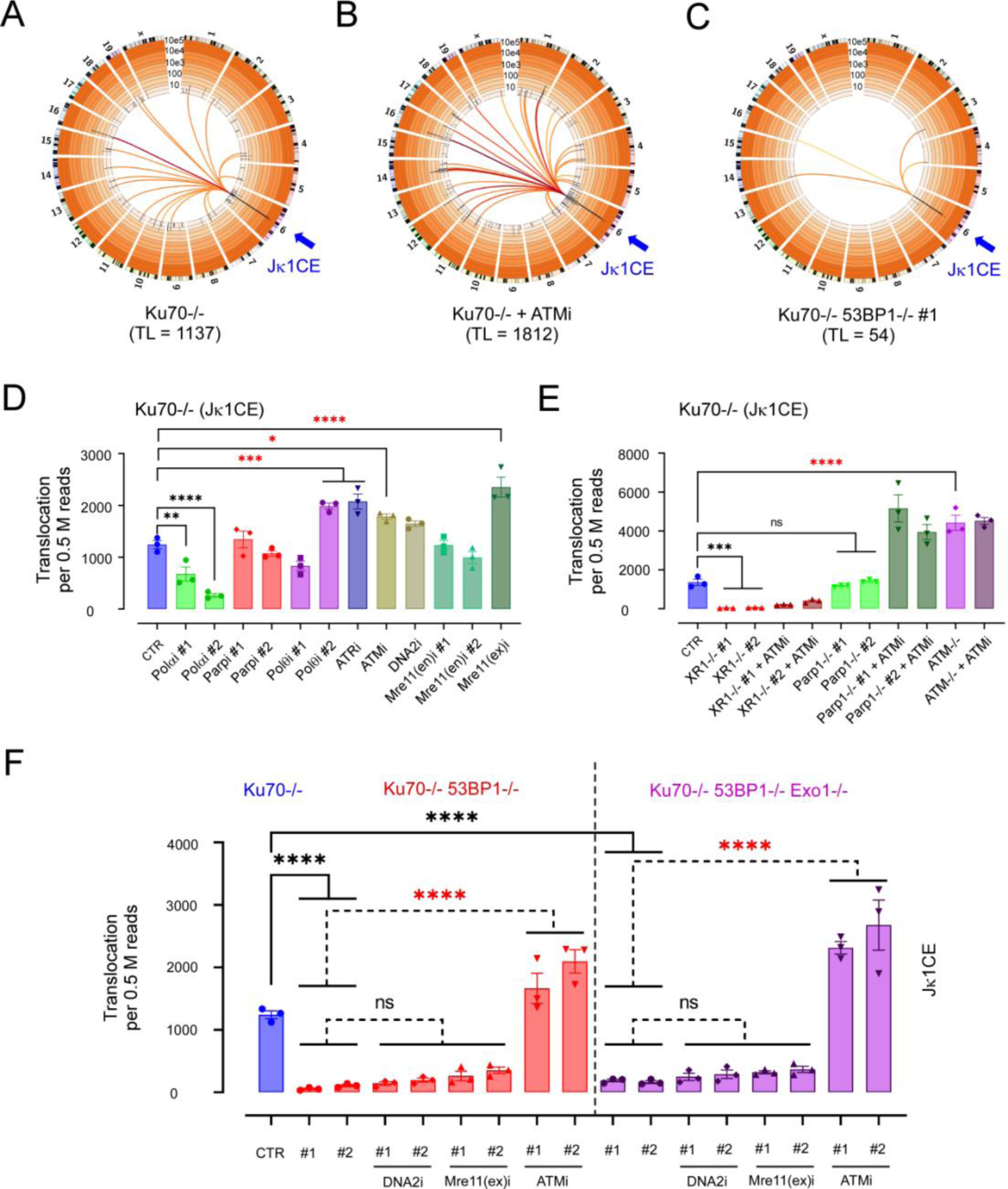
Compromised DDR affects A-EJ translocations. (**A-C**) Representative genome-wide plots of junctions joined with Jκ1CE in *Ku70^-/-^* (A), *Ku70^-/-^* + ATMi (B), *Ku70^-/-^ 53BP1^-/-^* #1 (C). (**D-F**) Relative translocation frequencies in *Ku70^-/-^* with/without indicated inhibitors, deletions, or both per 0.5 M (million) sequence read pairs. The significance between *Ku70^-/-^* (CTR) and indicated inhibitors, gene modification, or both were evaluated by one-way ANOVA plus post-test comparisons: * (p<0.05), ** (p<0.01), *** (p<0.001), **** (p<0.0001) and ns (no significance); N=3

The chemical inhibition and gene deletion effects on *Ku70^-/-^*translocation frequency revealed both gains and losses. ATM deletion or inhibition of ATR, Mre11(ex), or ATM (including with all gene deletion contexts) consistently increased translocations (Figs 7A-E, S20A-B), which, for the exception of Mre11(ex)i, corresponded with decreased V-J efficiencies (Figs. 1B-C, S4C-D). In contrast, deficiency of XRCC1 or 53BP1 recovered very few translocations (Figs. 7A-F, S20A-B, S21; Tables S3, S4) consistent with robust decreases in V-J efficiencies (Figs. 1C, 2B, S4D, S6B). Unlike with WT *vAbl* cells, Parpi or Parp1 deficiency in *Ku70^-/-^ vAbl* cells did not change the translocation level. This unexpected effect may be normalized due to a combined genome-wide DSB increase and a partially compromised A-EJ. Joint structures of *Ku70^-/-^* translocations resembled V-region joint structures, and similar patterns emerged for all other added DDR or A-EJ perturbations (Fig. S22A-D). Notably, ATMi treatment in all measurable *Ku70^-/-^* double knockout combinations generally doubled MMEJ utilization from the 20% baseline (Fig. S22C-D). However, the low translocation frequencies in the severely compromised backgrounds (i.e. *Ku70^-/-^Xrcc1^-/-^*and *Ku70^-/-^53bp1^-/-^*) precluded further comparison. In summary, the data reveal two distinct groups of genes that influence translocation generation: 1) upstream DDR genes that influence A-EJ synapsis partners, and 2) end joining capacity genes essential for the *bona fide* A-EJ pathway.

## Discussion

This study revealed Parp1 and XRCC1 as crucial A-EJ factors that drive Igκ V-J recombination and DSB repair in the absence of the Ku-initiated NHEJ. This mechanism is aided by the ATM-initiated DDR that stabilizes end-joining via 53BP1. On the one hand, disrupting the core A-EJ factor XRCC1 decreases both V-J and translocations, with residual joints comprising mostly of MHs to stabilize their end ligation. On the other hand, disrupting the DDR at different stages is varied, where key factor perturbations highlight defective end joining fidelity, efficiency, and/or synapsis functions, most of which are also either evident with or functionally compensated by NHEJ (41, 56). Thus, the short-range resection and repair that is associated with this A-EJ mechanism is an excellent model to understand the plasticity of the DDR, particularly from 53BP1-regulated and ATM-independent contexts to elucidate additional end-joining mechanisms (Fig. 8). Our data provide further insight into the regulation of this heir-apparent end-joining pathway.

**Fig. 8.**
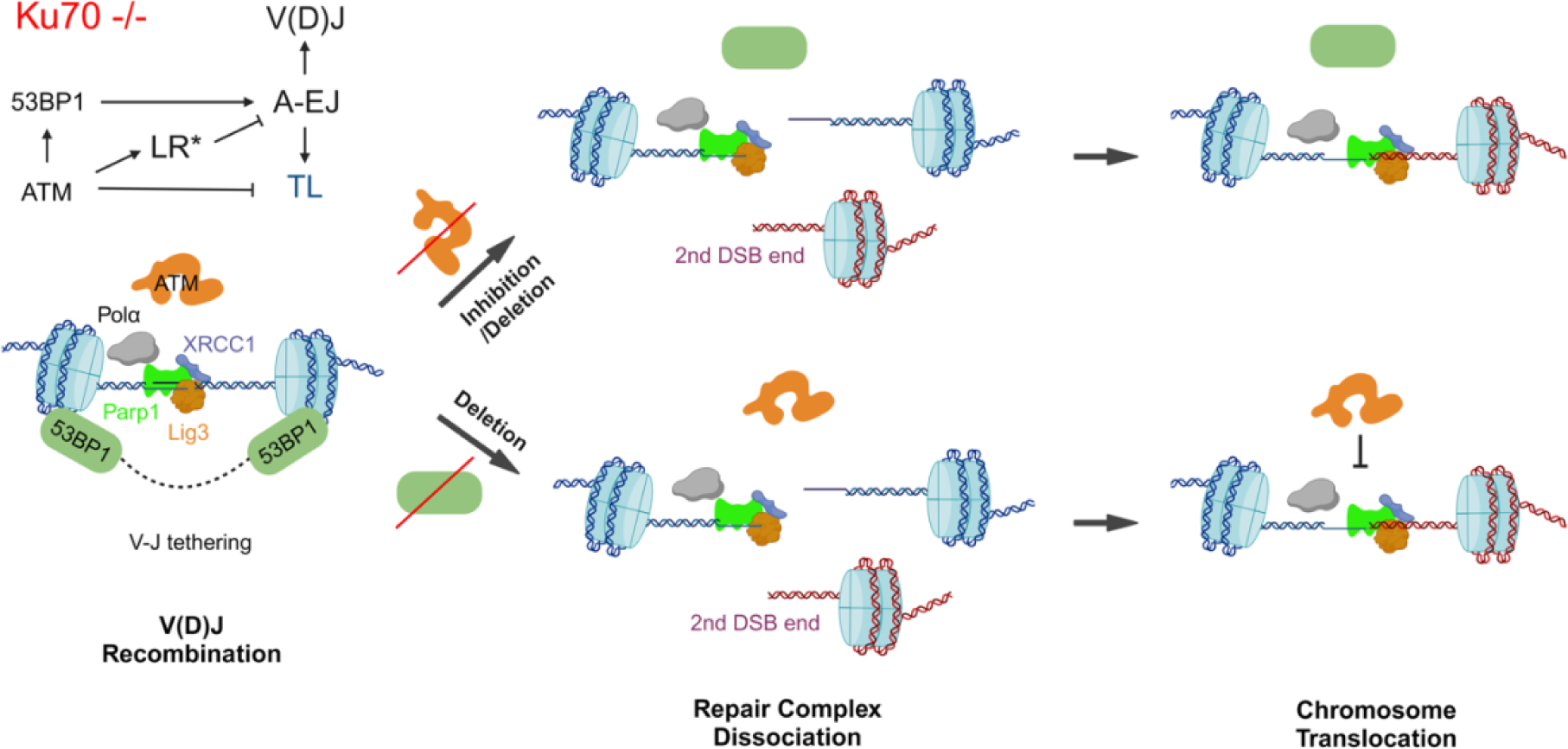
ATM and 53BP1 regulate A-EJ-mediated V(D)J recombination and chromosome translocation. Ku70 deficiency eliminates NHEJ and enables a *bona fide* A-EJ mechanism involving Parp1, XRCC1, and Lig3 to complete V(D)J recombination, initiated by the RAG1/2 endonuclease. A-EJ is supported by the ATM-mediated DDR which recruits 53BP1 to facilitate end salvage mechanisms (e.g. distal end tethering) and suppress the formation of translocations (TL). ATM also activates nucleases to promote long range resection (LR*) and suppress A-EJ when 53BP1 is absent or excluded from DDR recruitment. Thus, inhibiting or deleting ATM (top) increases translocations due to diminished end tethering support and nuclease activation, while deleting 53BP1 (bottom) decreases overall A-EJ capacity by dissociating the V-J tethered repair complex and enabling ATM-activated nucleases to suppress translocations.

Although its role(s) in A-EJ will need more clarity, poly(ADP)ribosylation contributed by Parp1 and Parp2 labels DNA end termini, histones, and DNA repair proteins (57, 58) and, therefore, could facilitate end access, tethering at the nucleosome level or even end bridging (59, 60). The likelihood of both genes operating in A-EJ is consistent with the partial loss in repair with Parp1 deficiency and a greater A-EJ defect with Parp inhibitors that preferentially trap Parp2 (61); however, functions in addition to A-EJ are likely the reason why dual deficiency of these genes is cell lethal in NHEJ-proficient *vAbl* cells (62). In this regard, Parp1/Parp2 functions in chromatin remodeling due to overlapping linker histone regulation with the DDR kinases (63, 64) may explain its role in supporting both DNA end joining mechanisms. Indeed, inhibiting ATM in *Ku70^-/-^Parp1^-/-^* cells had distinguishing phenotypes that were either synergistic (e.g. junction structures, translocations) or antagonistic (e.g. CE/SE ratio), with the latter involving an unscheduled release of SEs from the RAG1/2 post-cleavage complex. Thus, the Parp1/ATM redundancy to remove linker histones by a combination of phosphorylation, ubiquitination, or parylation, could be central to effectively joining DNA ends by the *bona fide* A-EJ mechanism.

Our data also implies Lig3 is the driving G0/G1 phase A-EJ ligase. Loss of its chromatin loader, XRCC1, significantly reduces its protein expression, whereas Lig1 protein expression is heavily downregulated at the onset of V(D)J recombination (41). The notable transition to more resected joints and increased MMEJ as a function of A-EJ loss, implicates XRCC1 as the central scaffolding factor necessary to form direct or limited MH joints. This model is consistent with prior studies demonstrating XRCC1 more rapidly accumulates at DNA strand breaks than the polymerase loader, PCNA (65, 66), and the DDR (67). In the context of base excision repair (BER), XRCC1 recruitment to damaged sites is significantly enhanced by Parp1 or Parp2 ADP ribosylation (68) and is a requisite component for polymerase beta (Polβ) recruitment (69) to enable limited fill-in activity. Curiously, polymerase lambda (Polλ) acts to “backup” Polβ excision repair functions (70). Both polymerases display some level of MMEJ activity for short 3’ overhangs (71), which, for Polλ, could generate short and long MHs that are characteristic of NHEJ and Polθ-mediated end joining, respectively (22). Therefore, we speculate that A-EJ may employ a similar BER polymerase preference, which may change with an increasing end destabilization burden.

With respect to how the DDR supports A-EJ, inactivation via ATM deletion or inhibition in *Ku70^-/-^*cells impaired A-EJ synapsis and enhanced MMEJ in all conditions, indicating a primary role to stabilize ends for ligation and preventing translocation (Fig. 8). The high functional overlap between the ATM perturbations for A-EJ is consistent with kinase-dead ATM versus deficiency having no overt difference in NHEJ-mediated V(D)J recombination despite kinase dead dominant effects to block topoisomerase I lesion resolution during replication as a putative causal mechanism for viability and cancer predisposition differences (72, 73). Therefore, the difference in V-J efficiency between inhibited (decreased) and deficient (no change) ATM could be due to the ability of ATR to compensate for ATM absence, given that inhibited ATR also negatively affected V(D)J recombination while increasing translocations; further study will be necessary to reveal unique and overlapping DDR kinase functions for A-EJ. However, the DDR’s most striking impact on A-EJ was 53BP1 deficiency, which robustly suppressed V-J efficiency and translocations while increasing MMEJ yet not altering resected joint distributions. These observations suggests that the 53BP1 chromatin domain provides end stability and alignment to facilitate blunt (direct) end joining, as evidenced by the numerous 53BP1-associated complexes to regulate end processing (31,35–37, 39), and presents a limited temporal window even for NHEJ to complete V(D)J recombination in its absence (74–76). Therefore, the residual A-EJ in the absence of 53BP1 likely reflects a kinetic component to DSB repair (i.e. fast ligation fraction). A corollary to the end stability provided by 53BP1 is that translocations are likely formed and perhaps synergized when persisting DSBs are fused into the same 53BP1 chromatin domain (77) as a necessary step to ensure stable DNA ends are repaired even at the cost of rearrangement.

Our study complements recent work described by the Sfeir and Ceccaldi groups (78, 79), which revealed a mitosis-specific DSB repair mechanism directed by the CDK1-activated polo-like kinase 1 (PLK1) to promote an ATM-dependent, but ATR-independent, MMEJ mechanism involving Polθ, TOPBP1, MDC1, the RAD9-RAD1-HUS1 DNA clamp complex, and its interacting nuclear orphan, RHINO. This mitotic DSB repair mechanism does not involve NHEJ or homologous recombination due to the negative regulation of 53BP1 and BRCA2 by the CDK1-PLK1 axis (78, 80–82). As cells enter G1-phase, NHEJ proficiency returns, enabling 53BP1 nuclear bodies to form due to unresolved mitotic DNA damage. Polθ nuclear bodies also form, but both are nearly mutually excluded from each other (78), suggesting that Polθ largely operates external to the 53BP1 domain in early G1 and is consistent with 53BP1 acting as a Polθ synthetic lethal partner (12, 83). Although further investigation will determine the repair and end processing hierarchy, we propose the *bona fide* A-EJ mechanism is functionally distinct from the Polθ-mediated A-EJ mechanism and speculate most of the MMEJ outcomes described here for A-EJ, elsewhere for *Lig4^-/-^* (4, 50), and in cycling cell contexts represent repair events beyond the regulation of 53BP1.

An added complication to this study is the regulation of the RAG1/2 post-cleavage synaptic complex by ATM and how these ends are handed off to repair pathways. Coding ends are first released from the RAG1/2 complex to complete repair by NHEJ, while the release of signal ends for repair occurs afterward. For the latter part, RAG2 phosphorylation by ATM suppresses the release of signal ends (53) to minimize bi-allelic cleavage (84). Therefore, premature signal end release as coding ends are processed would disrupt DNA-PK synapsis and generate more re-synapsed and ligated hybrid joints (41). In support of this model, our WT +ATMi *vAbl* data indicate signal ends are subjected to similar end processing as their fated coding-end partner, indicating re-synapsis occurs before most hairpins are opened. Furthermore, while the increased level of rejoined Jκ1 DSB ends occur in the absence of Ku70, ATMi in WT cells does not substantially promote this type of activity; rather, we find ∼10-fold increased inversional J-J coding joints to all Jκ DSBs in the Jκ recombination center which, given this high frequency relative to translocations to other antigen receptor loci (Table S3), suggests ATM regulates post cleavage activity of multiply-loaded RAG1/2 sites within a single Jκ recombination center. Therefore, with respect to A-EJ and the generation of substantially more hybrid joints, we propose the kinetics of signal end acquisition by A-EJ are faster than ATM recruitment (67) for most post-cleavage complexes, reflecting the modest increase in hybrid joints with additional ATM inhibition.

## Data Availability

Genome-wide junction maps and related sequencing data are on GEO under the ascension numbers GSE242952, GSE246239, and GSE263165.

## Acknowledgements

We thank John Tainer for the Mre11 inhibitors (PFM01, PFM03, PFM39) and the Stanford Cancer Institute core facilities. Some of the figure panels were generated by BioRender.com or adapted from the Integrative Genomics Viewer (igv.org).

## Author contributions

JW and RLF conceived the study, designed experiments, and interpreted the data. JW, CAS, and LVL performed experiments. MLB performed preliminary experiments and generated analysis scripts. JW drafted the manuscript. JW and RLF wrote the manuscript. RLF acquired funding and supervised the study.

