## Supplementary Figures for "ATM and 53BP1 regulate alternative end joining-mediated V(D)J recombination"

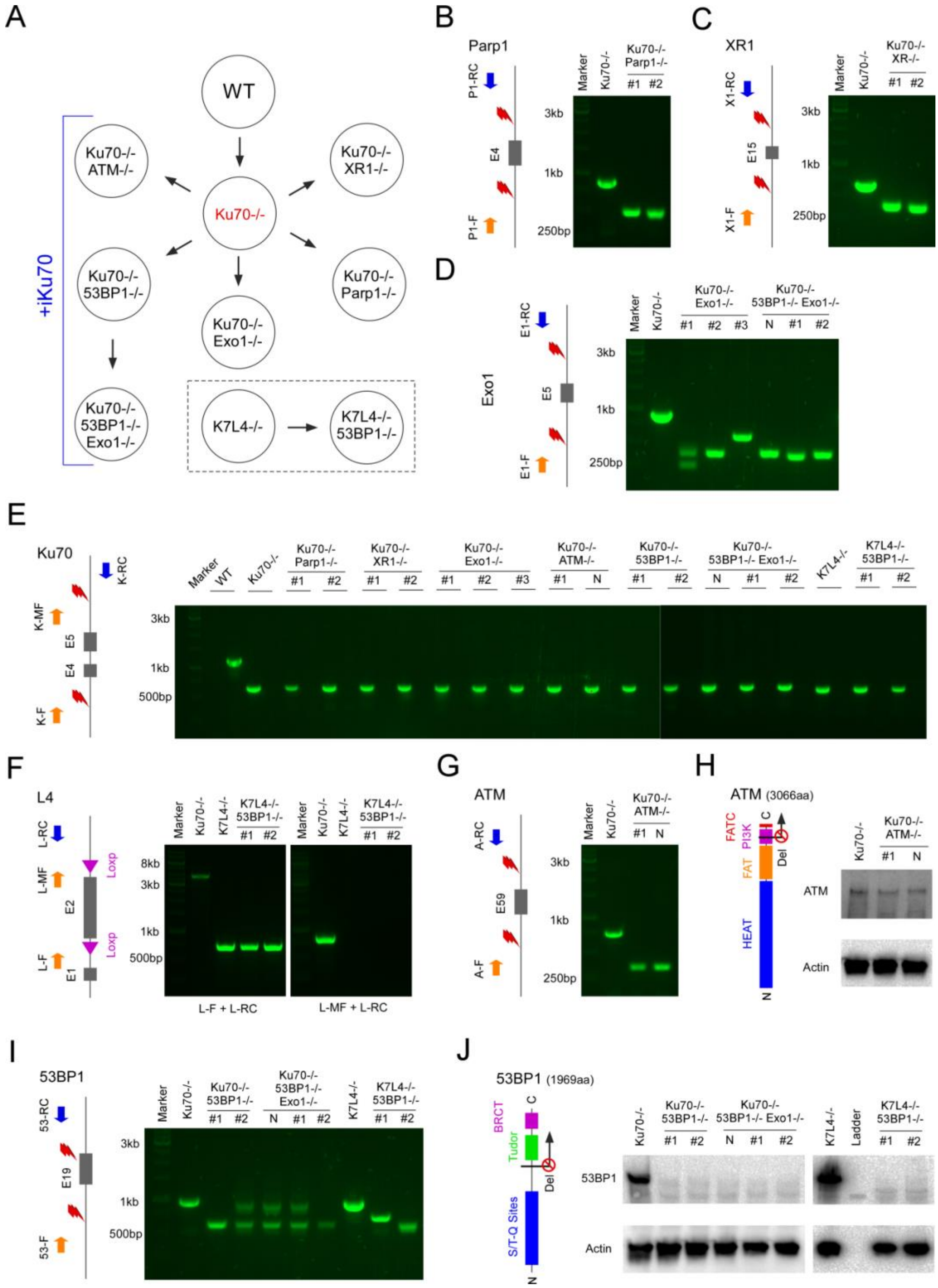


**Figure S1. Cell lines used for this study.** (**A**) The family tree of gene deleted *vAbl* cells. (**B**) The Exon 4 (E4) of Parp1 was deleted by Cas9 (lightning bolt symbols) and verified by genotyping using the indicated primers (P1-F and P1-RC). (**C-D**) Same as (B) except the Exon 15 (E15) of XRCC1 (XR1) and Exon 5 (E5) of Exo1 using indicated primers. (**E**) A larger DNA fragment (~3kb) containing Exon 4 (E4) and Exon 5 (E5) of Ku70 were deleted by Cas9. The corresponding cell lines and the WT control were verified by K-F plus K-RC (left) and K-MF plus K-RC (right) primer combinations. (**F**) Same as (E) except for Lig4 (L4) deletion by Cre-LoxP system and verified by indicated primer combinations. (**G-H**) Exon 59 (E59) of ATM, corresponding to the middle region of PI3K was deleted by Cas9 was verified by genotyping (G) that resulted in C-terminal loss due to frame shift further verified by western bloting (H). (**I-J**) Same as (G-H) except the Exon 19 (E19) deletion of 53BP1 and loss of expression of whole protein.

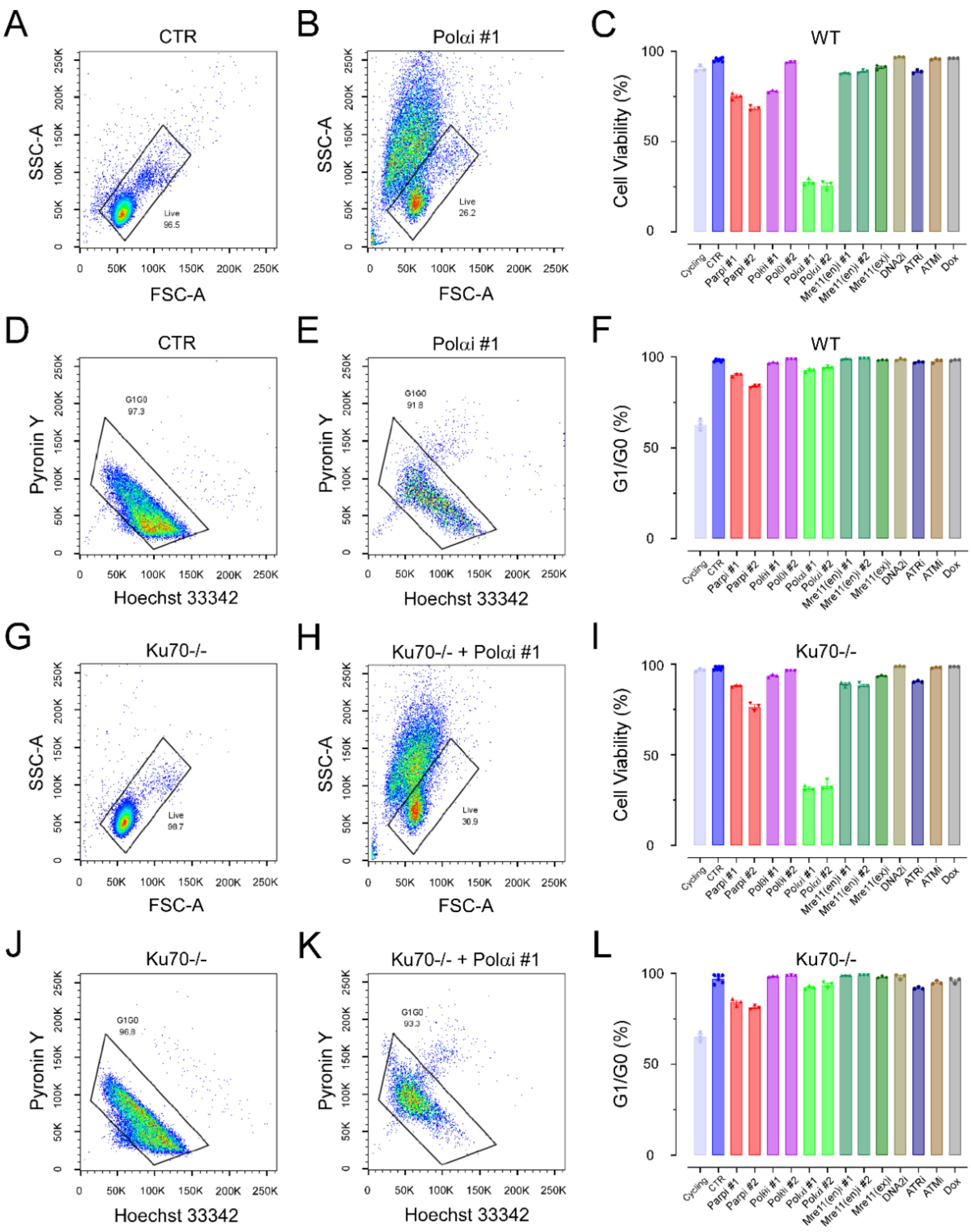


**Figure S2. Cell death and cell cycle analysis by flow cytometry.** (**A-B**) Representative Figures of cell viability analysis in the control (A) and Polαi-treated conditions (B). Dead cells (ungated) and live cells (gated) displayed distinct clusters in the FSC-A and SSC-A channels. (**C**) The percentages of live *vAbl* WT cells with/without indicated compounds used for this study. (**D-E**) Representative Figures of cell cycle analysis in the control (D) and Polαi-treated conditions (E), with Hoechst 33342 staining the content of genomic DNA while Pyronin Y staining the RNA. The gating indicates the population of G1/G0 cells. (**F**) The G1/G0 populations in the condition with/without indicated compounds were shown in the bars. (**G-L**) Same as (A-F) except for the *Ku70*$^{-/-}$.

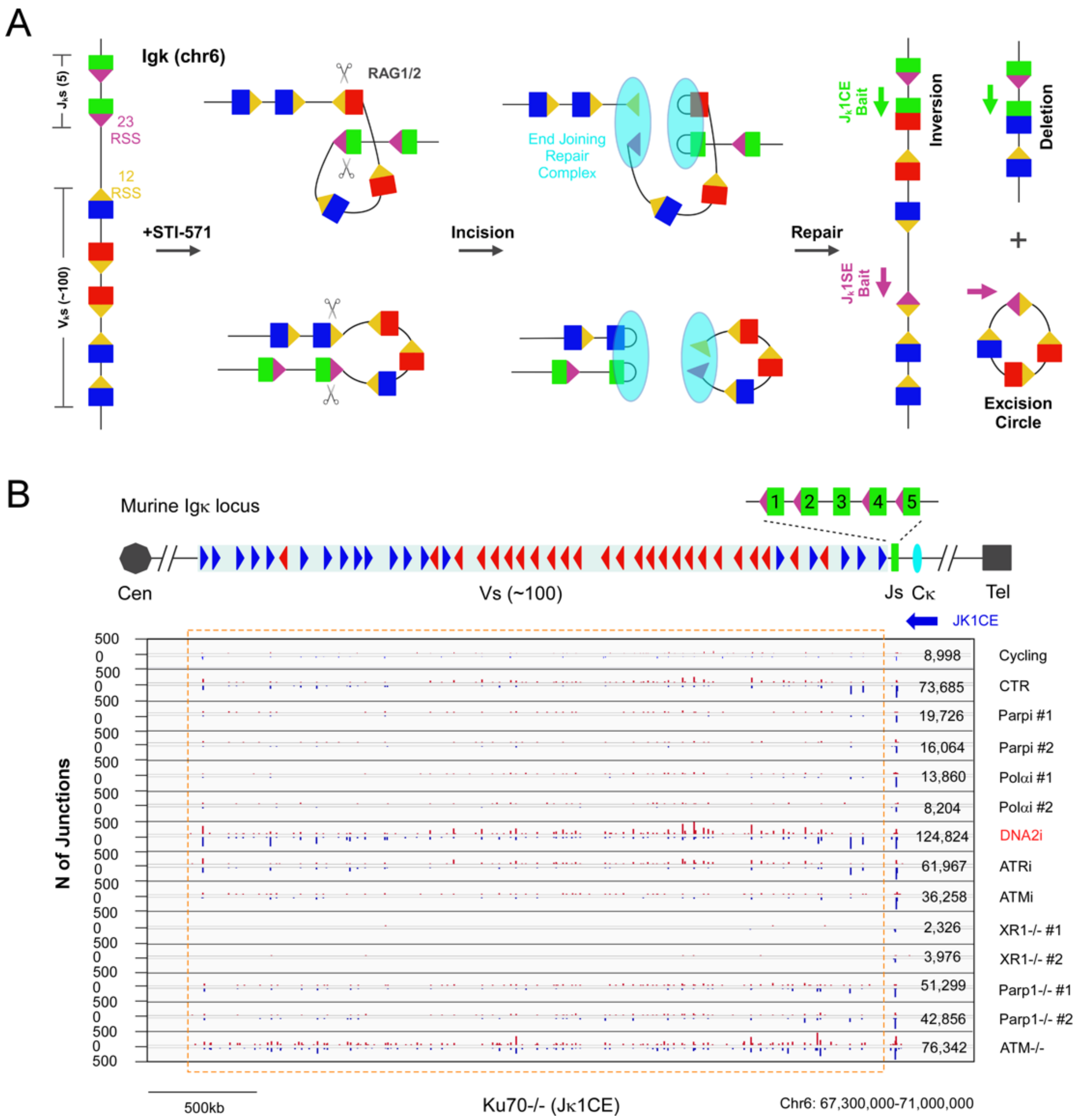


**Figure S3. Representative landscapes of V-J junctions captured by Jκ1CE.** (**A**) The Igκ antigen receptor locus contains inversional (red boxed) and deletional (blue boxes) V gene segments with respect to the J gene segments (green boxes). Recombination of V and J gene segment regions in *vAbl* cells is initiated by RAG1/2 endonuclease V-J pairing and incision (scissors) between each gene segment and its associated recombination signal sequences (RSS) distinguished by non-conserved internal 12 or 23 bp spacer sequence (triangles), followed by NHEJ to generate inversional and deletional coding joints and their complementary signal joints which include the other half of the inversion and the interstitial region forming an excision circle, respectively. See Figure S5 for WT recombination outcomes. A-EJ (*Ku70*$^{-/-}$) promotes substantially more hybrid (coding/signal) joints with outcomes between V and J regions that are depicted in Figs. 1 and S4. (**B**) Representative plots display Vκ region junctions (orange dashed box) that recombined to the Jκ1 coding end (Jκ1CE) in *vAbl Ku70*$^{-/-}$ cells without STI-571 treatment (cycling), and with STI-571 treatment combined with DMSO (CTR), inhibitors (Parp1i #1/2, Polθi #1, Polαi #1/2, DNA2i, ATRi and ATMi) or gene deletions (XRCC1 - XR1$^{-/-}$ #1/2, Parp1$^{-/-}$ #1/2 and ATM$^{-/-}$). The total Vκ region junctions (orange rectangular) in either plus (red) or minus (blue) DNA strand orientations are indicated. Scale bar is 500kb.

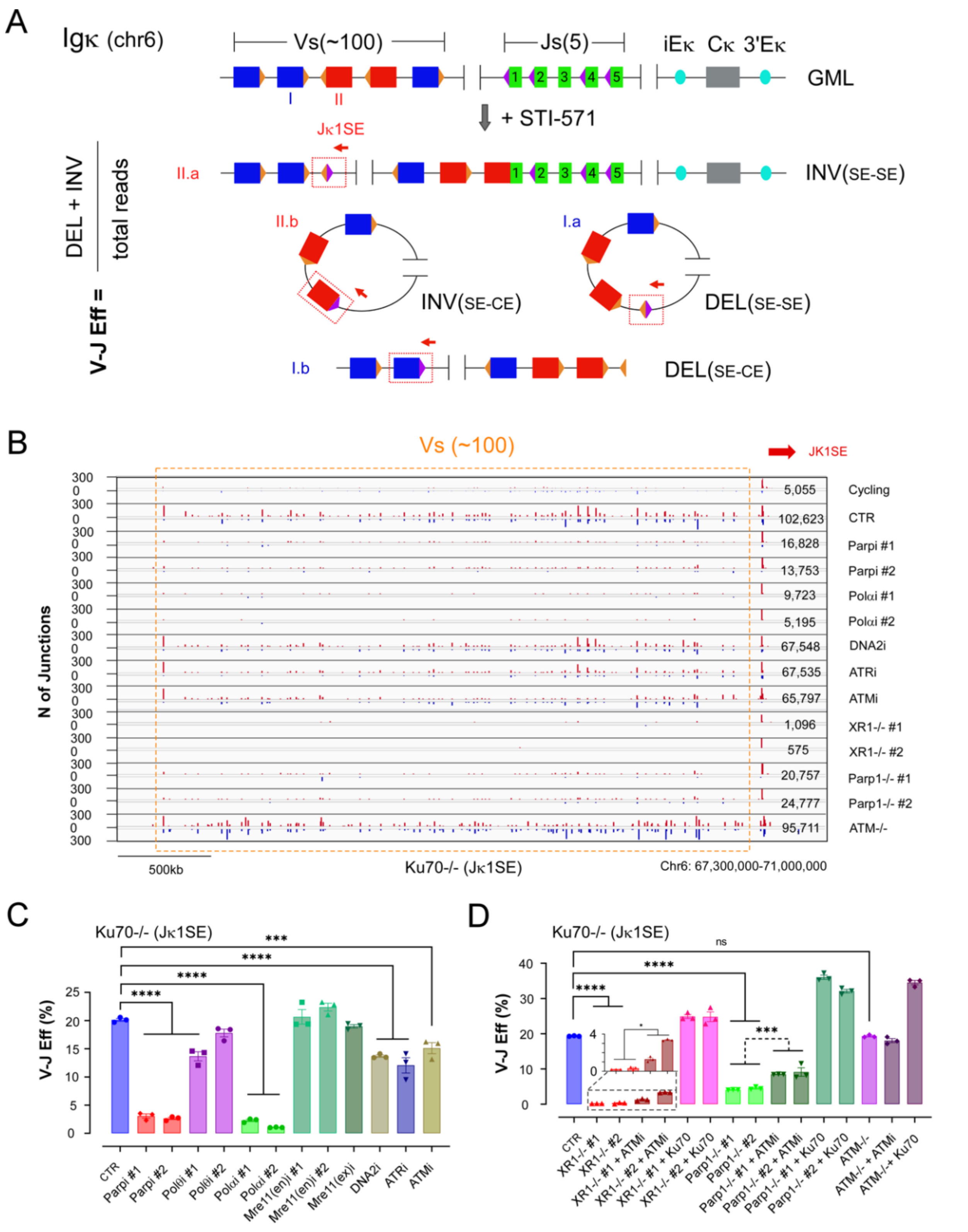

**Figure S4. Characterizing the genes that affect Ku70-independent V-J signal end recombination efficiency.** (**A**) Representative Vκ region junctions that recombined to the Jκ1 signal end (Jκ1SE) in *vAbl Ku70*$^{-/-}$ cells from inversion-oriented Vκ gene segments (INV, red; example: II) forming inversion signal joints (SE-SE; II.a) or excision circle hybrid joints (SE-CE; II.b) and deletion-oriented Vκ gene segments (DEL, blue; example: I) forming excision circle signal joints (SE-SE; I.a) or inversional hybrid joints (SE-CE; I.b) captured by the Jκ1SE bait primer (red arrow). The V-J Eff was calculated by the sum of INV and DEL divided by the total sequence reads. GML = germline configuration. (**B**) Representative plots display junctions that recombined to Jκ1SE in *vAbl Ku70*$^{-/-}$ cells under cycling or STI-571 conditions (everything else) with added inhibitors (Parpi #1/2, Polθi #1/2, Polαi #1/2, DNA2i, ATRi and ATMi) or gene deletions (XRCC1 - *XR1*$^{-/-}$ #1/2, *Parp1*$^{-/-}$ #1/2 and *Atm*$^{-/-}$). The total numbers of junctions in Igκ V region (orange rectangle) in either plus (red) or minus (blue) DNA strand orientations are indicated. Scale bar is 500kb. (**C**) V-J recombination efficiency of *vAbl Ku70*$^{-/-}$ cells with or without inhibitors are indicated. (**D**) V-J recombination efficiency of *vAbl Ku70*$^{-/-}$ cells with gene deletions or with added ATMi or Ku70 expression are indicated. Differences in (C-D) were evaluated by One-way ANOVA plus post-comparison, with *** ($p<0.001$), **** ($p<0.0001$) and ns (no significance). Differences in the ATMi conditional subgroups *XR1*$^{-/-}$ and *Parp1*$^{-/-}$ were evaluated by Two-way ANOVA plus post-comparison with * ($p<0.05$), *** ($p<0.001$). All the experiments were independently repeated three times.

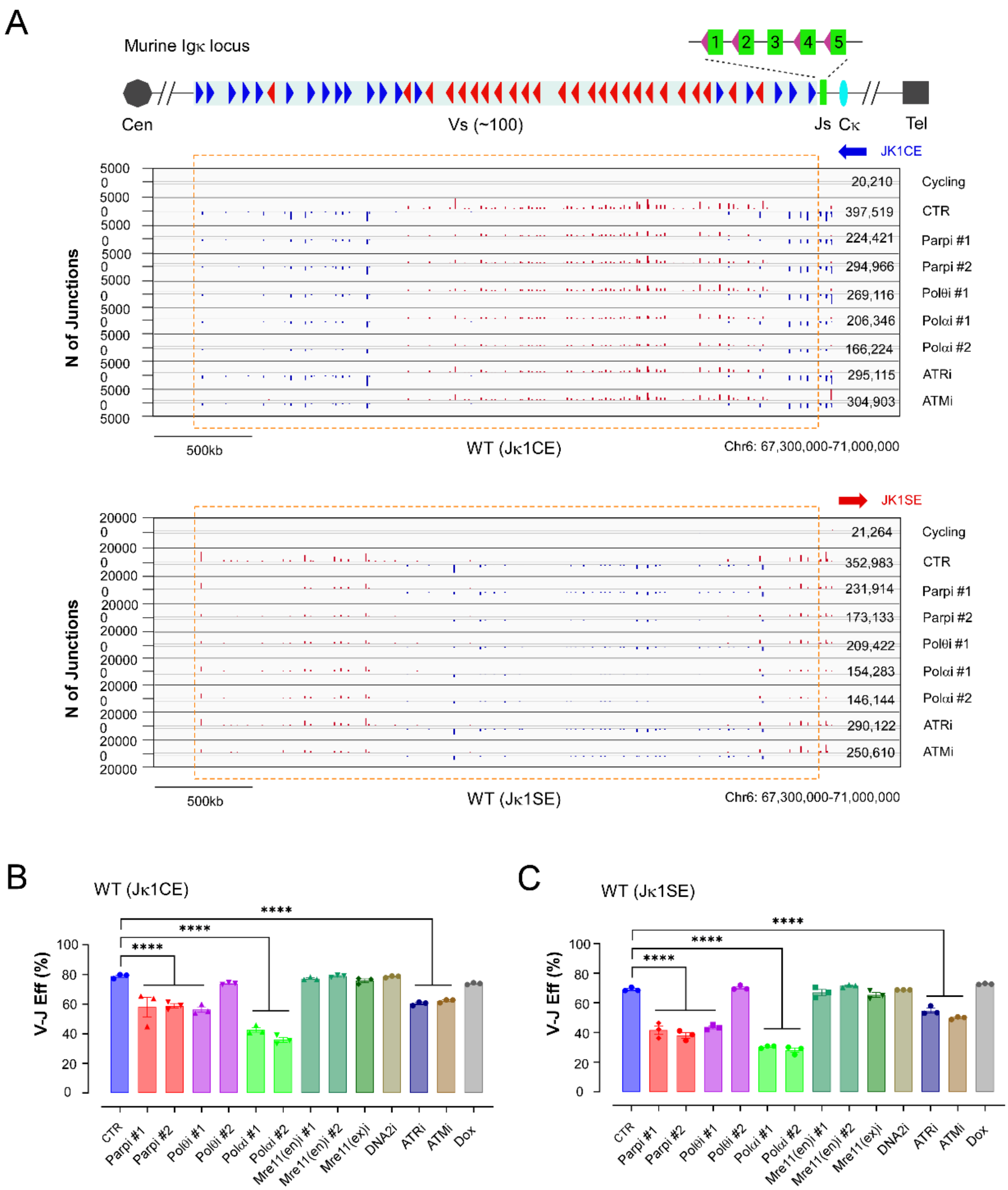


**Figure S5. Characterizing the genes that affect V-J recombination efficiency in *vAbl WT* cells.** (**A**) Murine Igκ antigen locus consists of ~100 V gene segments (blue and red triangles depicting their associated recombination signal sequence), five J gene segments (green) and constant gene backbone Cκ (cyan). The representative plots of junctions that recombined to Jκ1CE and Jκ1SE baits in *vAbl* WT cycling cells or with the treatment of STI-571 coupled with inhibitions DMSO (CTR), Parpi #1/2, Polθi #1/2, Polαi #1/2, ATRi and ATMi. The total junction numbers in the Vκ region (orange rectangle) are indicated. Scale bar is 500kb. (**B**) V-J recombination efficiency derived from the Jκ1CE bait with or without inhibitors, including CTR (DMSO), Parpi #1/2, Polθi #1/2, Polαi #1/2, Mre11(en)i #1/2, Mre11(ex)i, DNA2i, ATRi, ATMi and Doxycline (Dox). (**C**) Same as (B) but for Jκ1SE bait. The differences in (B-C) were obtained by One-way ANOVA plus post-comparison, with **** (p<0.0001). All the experiments were independently repeated three times.

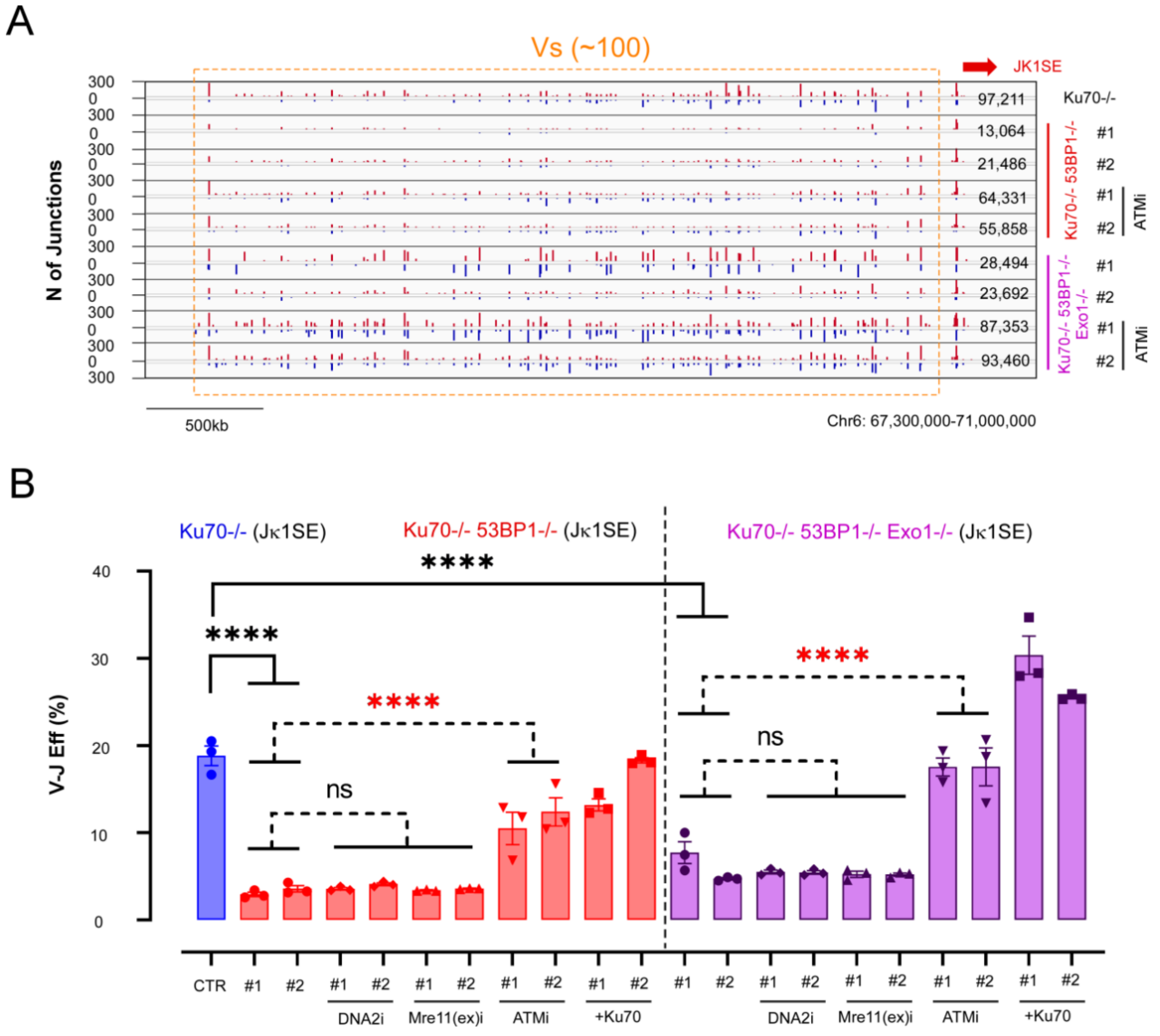


**Figure S6. ATMi partially restores V-J signal end recombination in *Ku70⁻/⁻* cells caused by 53BP1 depletion.** (**A**) Representative plots of junctions that recombined to Jκ1SE bait in *Ku70⁻/⁻*, *Ku70⁻/⁻53bp1⁻/⁻* #1/2 and *Ku70⁻/⁻53bp1⁻/⁻ Exo1⁻/⁻*, with or without ATM inhibition. (**B**) The V-J recombination efficiency in *Ku70⁻/⁻*, *Ku70⁻/⁻53bp1⁻/⁻* #1/2 and *Ku70⁻/⁻53bp1⁻/⁻Exo1⁻/⁻*, with or without DNA2, Mre11(ex) and ATM inhibitors, or Ku70 rescue expression. The difference between *Ku70⁻/⁻* (CTR) and rest treatment conditions are evaluated by One-way ANOVA plus post-comparison, and the differences within *Ku70⁻/⁻53bp1⁻/⁻* (red bars) and *Ku70⁻/⁻53bp1⁻/⁻Exo1⁻/⁻* (magenta bars) were evaluated by two-way ANOVA plus post-comparison, with **** ($p<0.0001$) and ns (no significance). All the experiments were independently repeated three times.

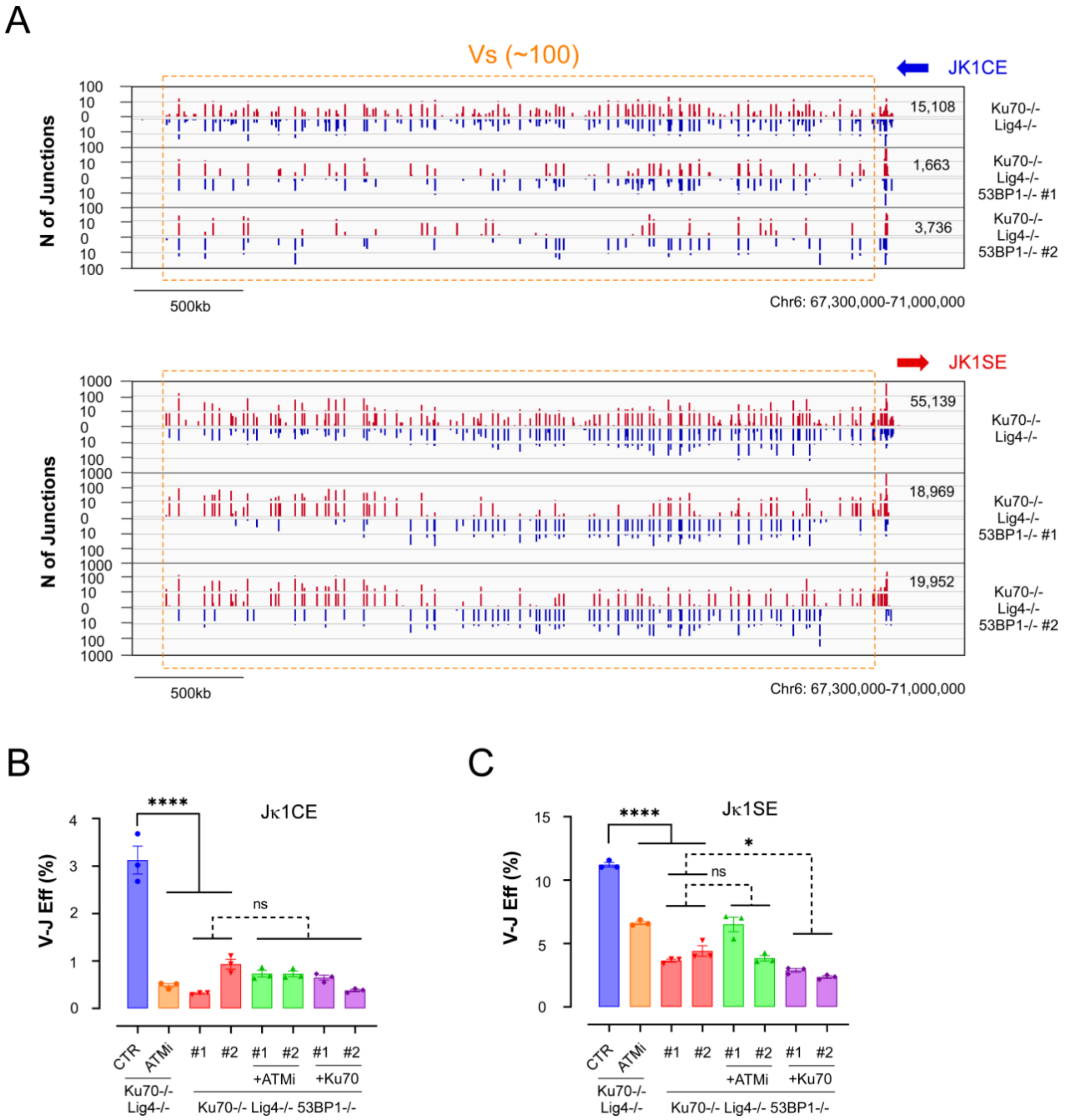


**Figure S7. ATMi does not restore V-J recombination in *Ku70$^{-/-}$Lig4$^{-/-}$* cells caused by 53BP1 depletion.** (**A**) Representative plots of junctions that recombined to Jκ1CE and Jκ1SE baits, respectively, in *Ku70$^{-/-}$Lig4$^{-/-}$* and *Ku70$^{-/-}$Lig4$^{-/-}$ 53bp1$^{-/-}$* #1/2. (**B-C**) V-J recombination efficiency in Ku70$^{-/-}$ Lig4$^{-/-}$ and *Ku70$^{-/-}$ Lig4$^{-/-}$ 53bp1$^{-/-}$* #1/2, with or without ATM inhibition or Ku70 rescue expression using Jκ1CE and Jκ1SE, respectively. The difference between *Ku70$^{-/-}$Lig4$^{-/-}$* and treatment conditions were evaluated by one-way ANOVA plus post-comparison, and the differences within *Ku70$^{-/-}$ Lig4$^{-/-}$ 53bp1$^{-/-}$* (red bars) and further treated conditions were evaluated by Two-way ANOVA plus post-comparison, with * (p<0.05), **** (p<0.0001) and ns (no significance). All the experiments were independently repeated three times.

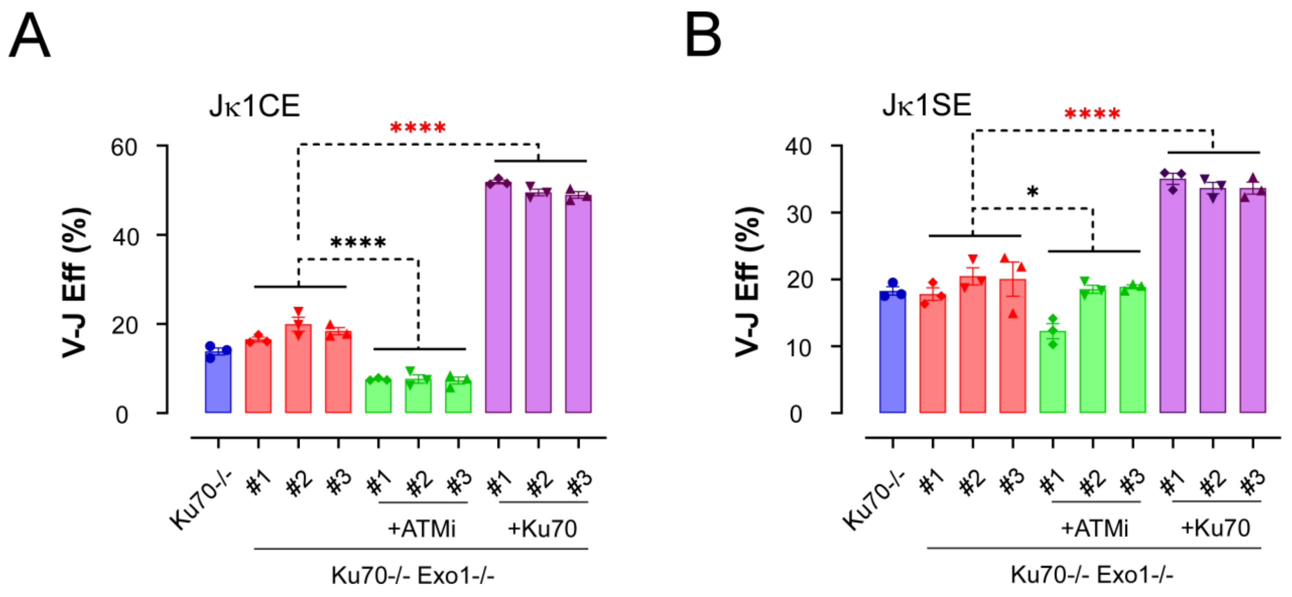


**Figure S8. Exo1 has little impact on A-EJ mediated V-J recombination.** (**A**) V-J coding end recombination efficiency in *Ku70*$^{-/-}$, *Ku70*$^{-/-}$ *Exo1*$^{-/-}$ #1/2/3 with or without ATM inhibitor or Ku70 rescue expression. The changes between *Ku70*$^{-/-}$ *Exo1*$^{-/-}$ and indicated treatment conditions were evaluated by two-way ANOVA plus post-comparison with **** ($p<0.0001$). (**B**) Same as (A) except with the signal end bait: * ($p<0.05$) and **** ($p<0.0001$). All the experiments were independently repeated three times.

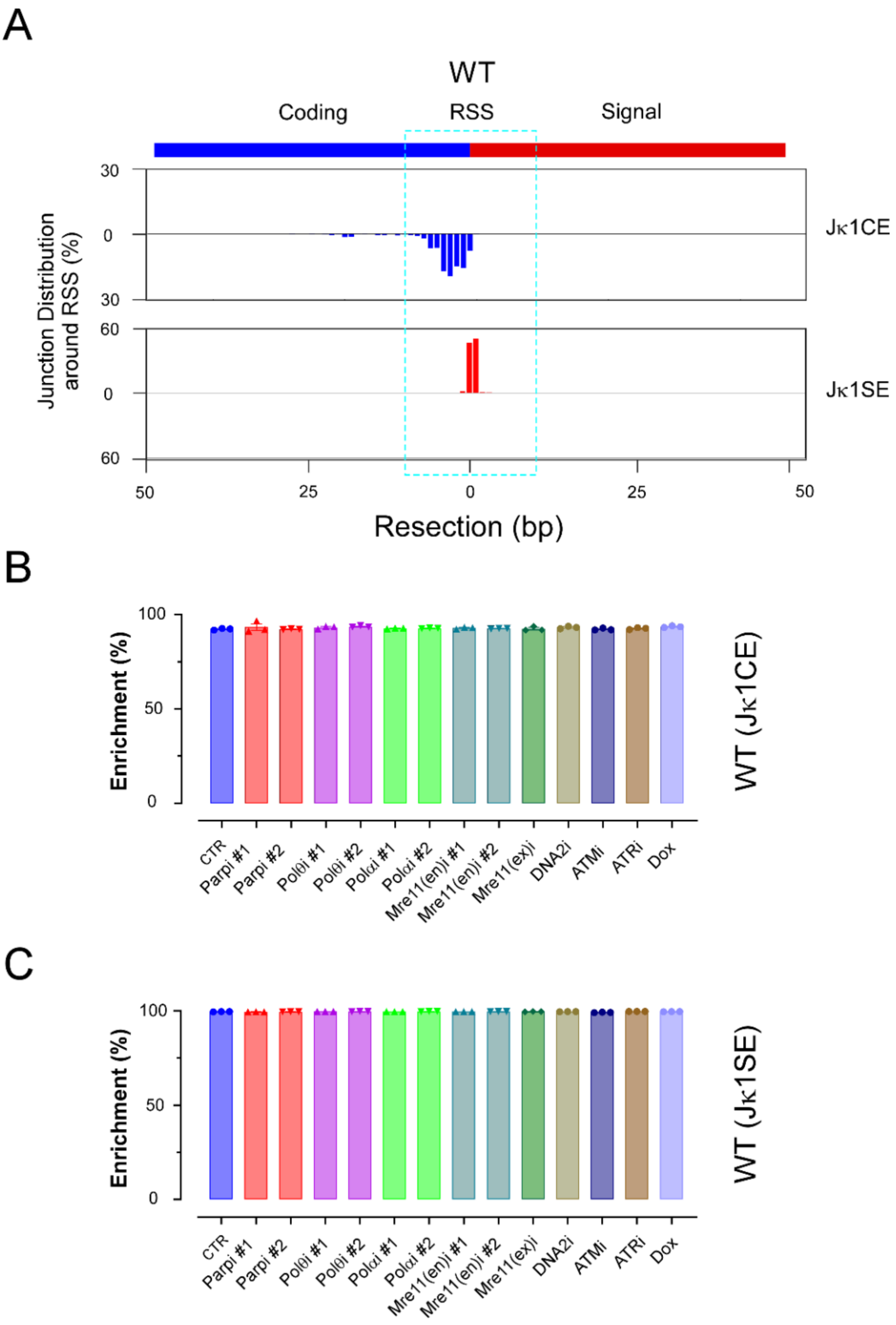


**Figure S9. Identifying genes regulating end resection in WT *vAbl* cells.** (**A**) The 100+ Vκ recombined junction distributions are collapsed onto an absolute position map with the left side representing all Vκ coding and the right side representing all Vκ signal positions with respect to RAG1/2 incision (zero point). The limited resection window of enrichment (±10 bps around DSB) is indicated (dashed cyan rectangle). (**B-C**) The percentage of junctions that enriched in the 10bp resection window was used to discern changes to resection. The values of enrichment in WT combined with indicated inhibitors captured by Jκ1CE (B) and Jκ1SE (C) baits were shown in bar graphs. All the experiments were independently repeated three times.

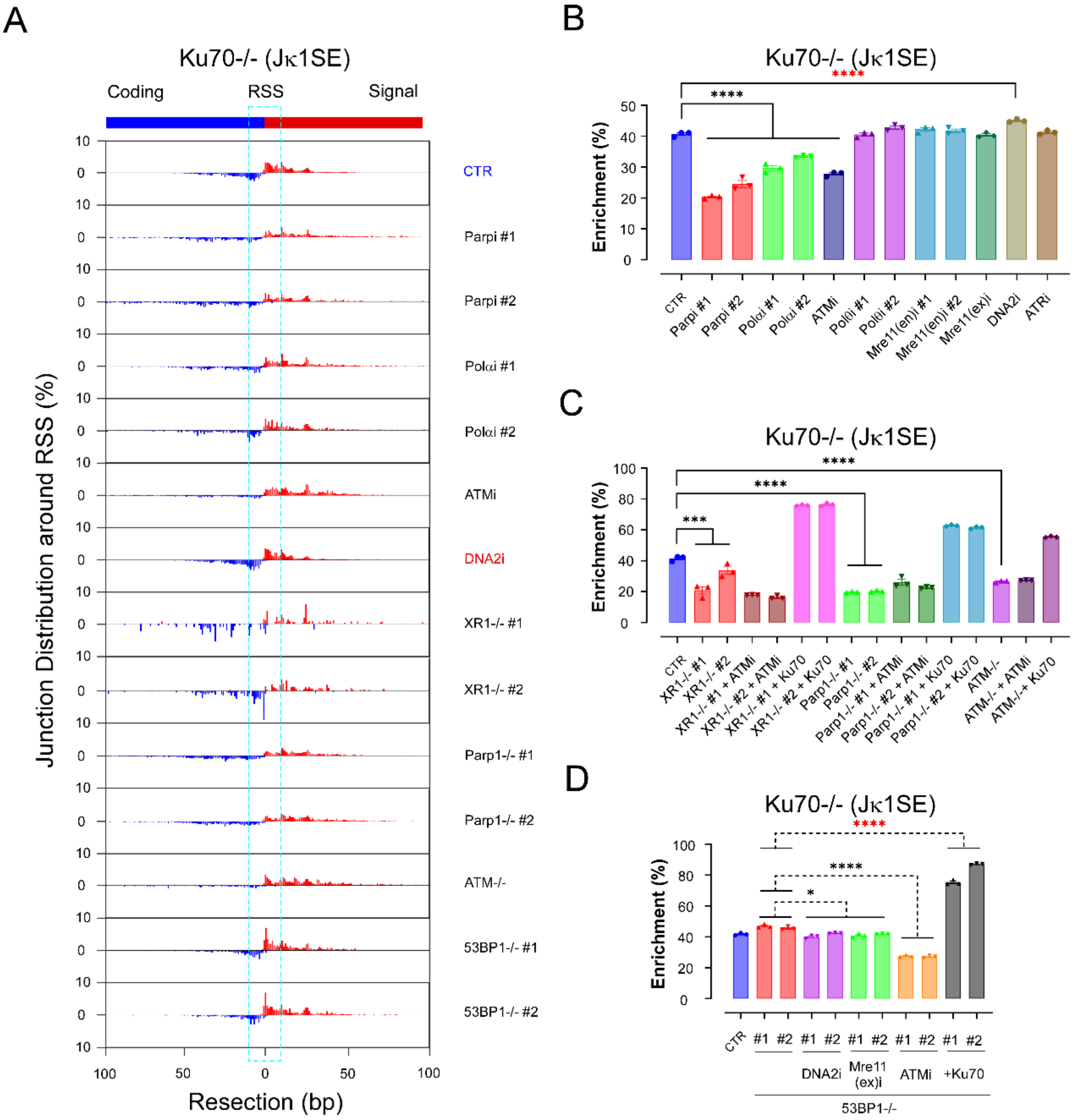


**Figure S10. Identifying genes that regulate prey resected joint distributions from the Jκ1SE bait.** (A) In the absence of Ku70, the coding and signal ends of V segments joined with Jκ1SE in a resection window of 200 bps (±100 bps around RSS site). The representative plots showing the degrees of resection was restricted (DNA2i treatment), extended (Polα inhibition; XRCC1 deletion; Parp1 and ATM inhibition or deletion), or no significant change (53BP1 deletion). (B-D) The percentage of junctions that enriched within ±10 bps of the RAG1/2 DSB (dashed cyan rectangle in (A)) was used measure resection changes. The values of enrichment in *Ku70*$^{-/-}$ combined with indicated inhibitors, gene modification or both were evaluated by one-way ANOVA plus post comparison (B-C) and two-way ANOVA plus post comparison (D), with * (p<0.05), ** (p<0.01) and **** (p<0.0001). All the experiments were independently repeated three times.

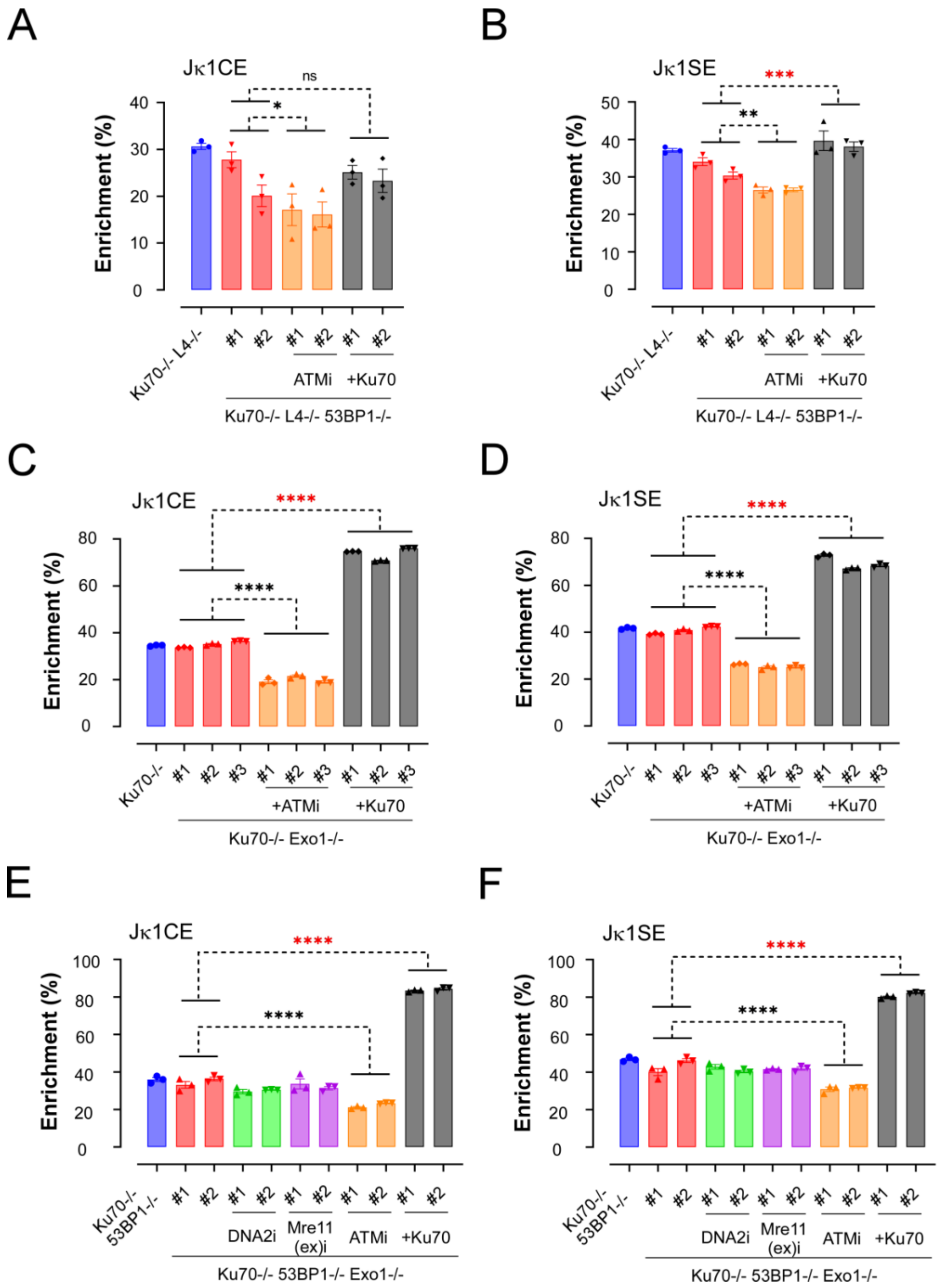


**Figure S11. Exo1 and 53BP1 do not impact resected V region joint patterns.** (**A-B**) The percentages of junctions that enriched within ±10 bps of Vκ DSB sites for *Ku70*$^{-/-}$*Lig4*$^{-/-}$, *Ku70*$^{-/-}$*Lig4*$^{-/-}$*53bp1*$^{-/-}$ *vAbl* cells with or without ATMi or Ku70 rescue expression from Jκ1CE (A) or Jκ1SE (B) baits, respectively. (**E-F**) Similar as (A-B) but for additional Exo1 knockout on top of *Ku70*$^{-/-}$. (**E-F**) Similar as (A-B) but for additional Exo1 knockout on top of *Ku70*$^{-/-}$*53bp1*$^{-/-}$ with extra inhibitors, DNA2i and Mre11(ex)i, tested. The changes within each group were evaluated by two-way ANOVA plus post comparison with * (p<0.05), ** (p<0.01), *** (p<0.001), **** (p<0.0001) and ns (no significance). All the experiments were independently repeated three times.

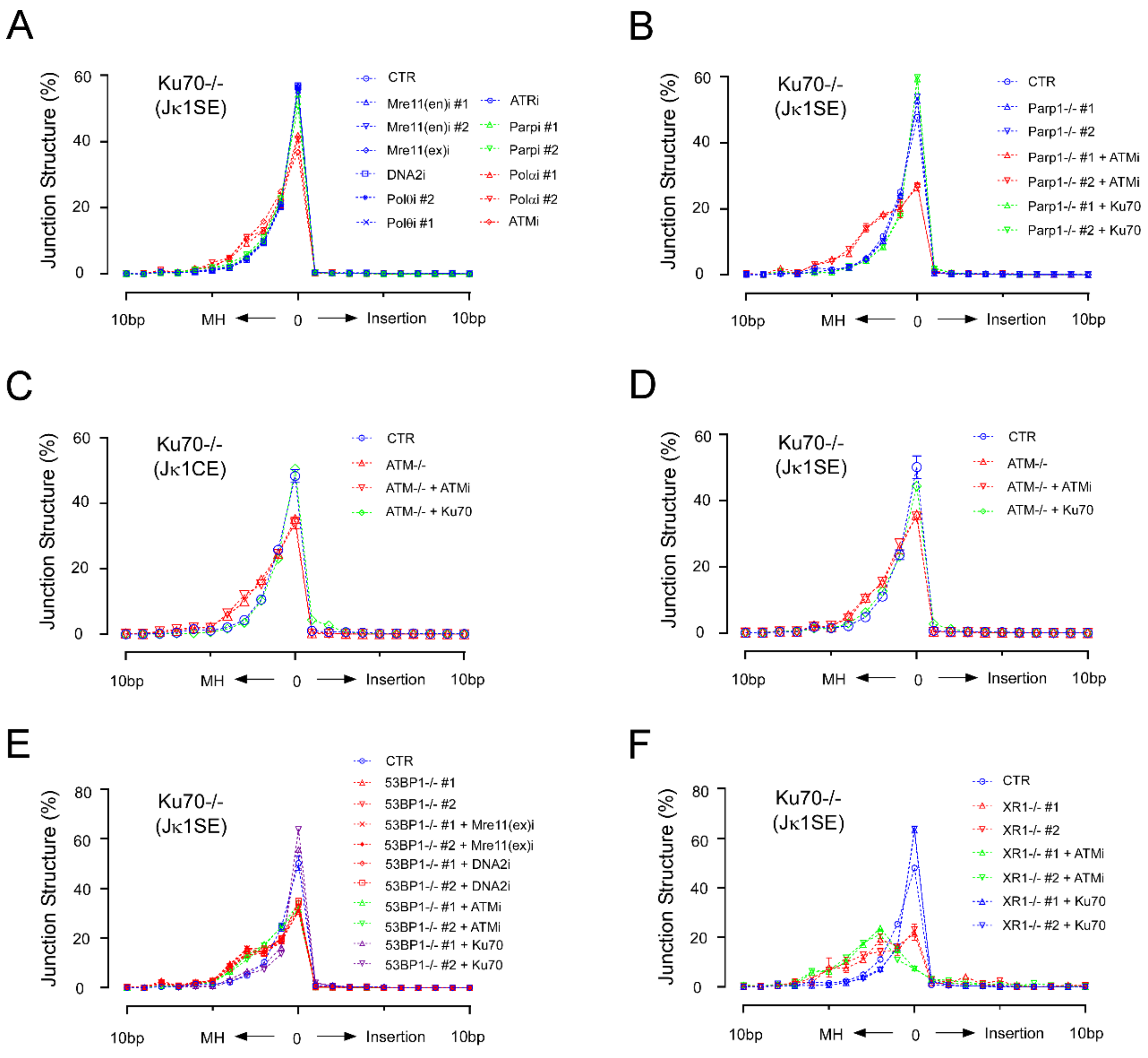


**Figure S12. Polα, ATM, 53BP1, Parp1, and XRCC1 compensate for the microhomology dependency of V region recombined joints.** (**A**) The distribution of junction structures including microhomology, direct repair and insertion in the conditions of *Ku70*$^{-/-}$ with/without indicated inhibitors using Jκ1SE as bait, with no detectable pattern change (blue), marginal change (Parpi #1/2, green) and significant change (Polαi #1/2 and ATMi). (**B-F**) Same as in (A) but for Parp1, ATM, 53BP1 and XRCC1 deletion with/without indicated inhibitors or Ku70 rescue expression, respectively, except Jκ1CE was used in (C). All the experiments were independently repeated three times.

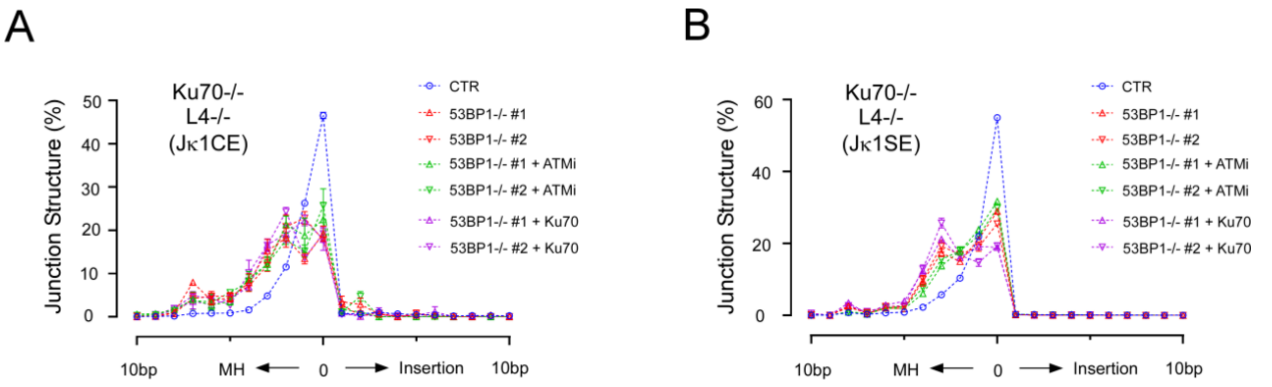


**Figure S13. *53bp1$^{-/-}$* dramatically enhances the MH usage in the absence of Ku70 and Lig4 (L4).** (**A**) The distribution of junction structures, including microhomology, direct repair and insertion, in *Ku70$^{-/-}$Lig4$^{-/-}$* (CTR) and *Ku70$^{-/-}$Lig4$^{-/-}$53bp1$^{-/-}$* #1/2 *vAbl* cells with/without ATMi and Ku70 rescue expression, obtained by Jκ1CE. (**B**) Same as (A) but for Jκ1SE. All the experiments were independently repeated three times.

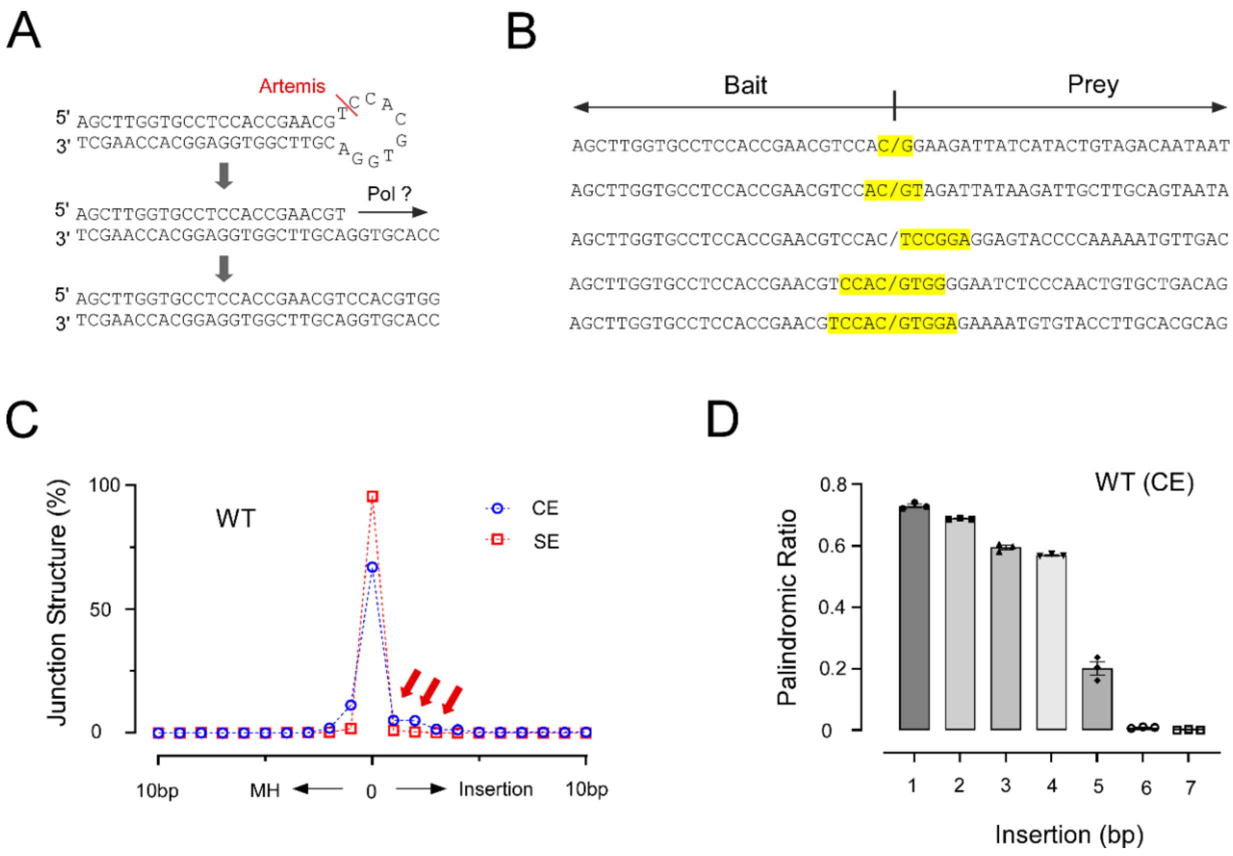


**Figure S14. Palindromic insertions from hairpin opening and fill-in are the dominant coding end insertions in WT *vAbl* cells.** (**A**) The mechanism of palindromic insertion generation, where hairpin of the coding end was opened by Artemis, followed by unknown polymerase. (**B**) Representative palindromic insertions (highlighted) obtained from the Jκ1CE bait; prey indicates the contribution from V genes. (**C**) Insertions (red arrows) were limited to the coding end. (**D**) Distribution of palindromic coding end insertion lengths relative to random insertion.

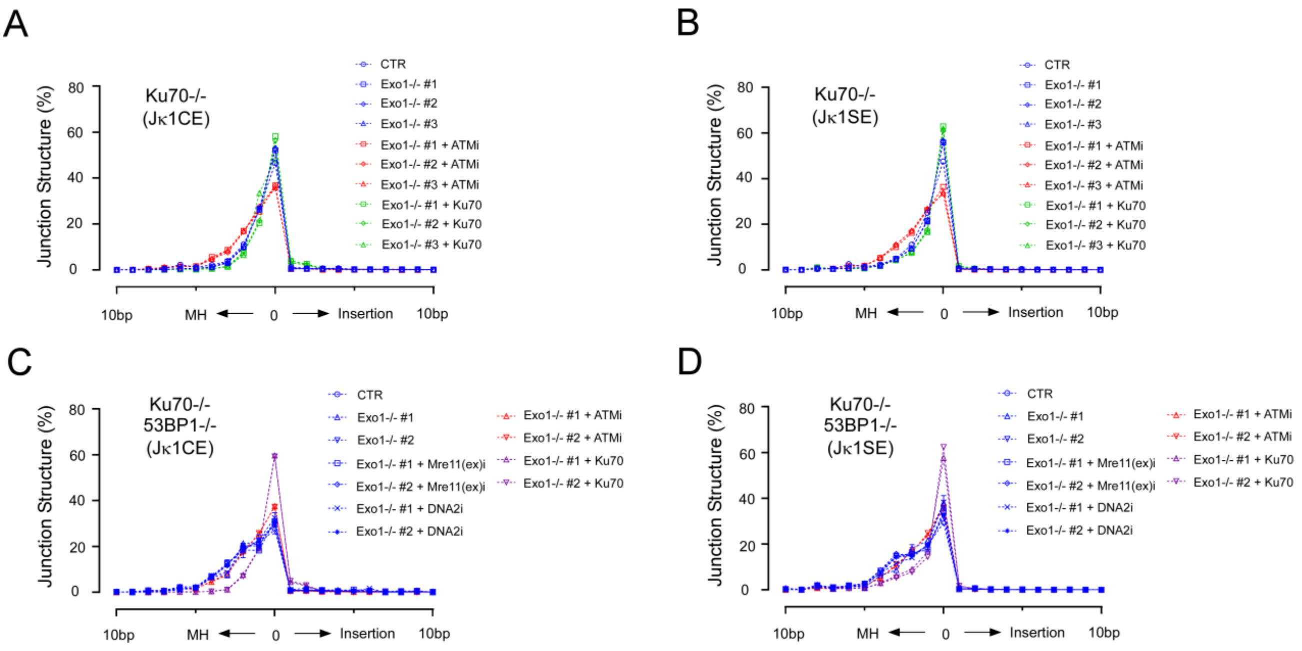


**Figure S15. Exo1 deficiency did not change the junction structures of V region recombinants by A-EJ.** (**A-B**) The distribution of junction structures for in the conditions of *Ku70*$^{-/-}$*Exo1*$^{-/-}$ (#1/2/3) with/without ATMi or Ku70 rescue expression. (**C-D**) Same as (A-B) but for *Ku70*$^{-/-}$*53bp1*$^{-/-}$ parental control and additional Exo1 deletion with/without ATMi or Ku70 rescue expression. All the experiments were independently repeated three times.

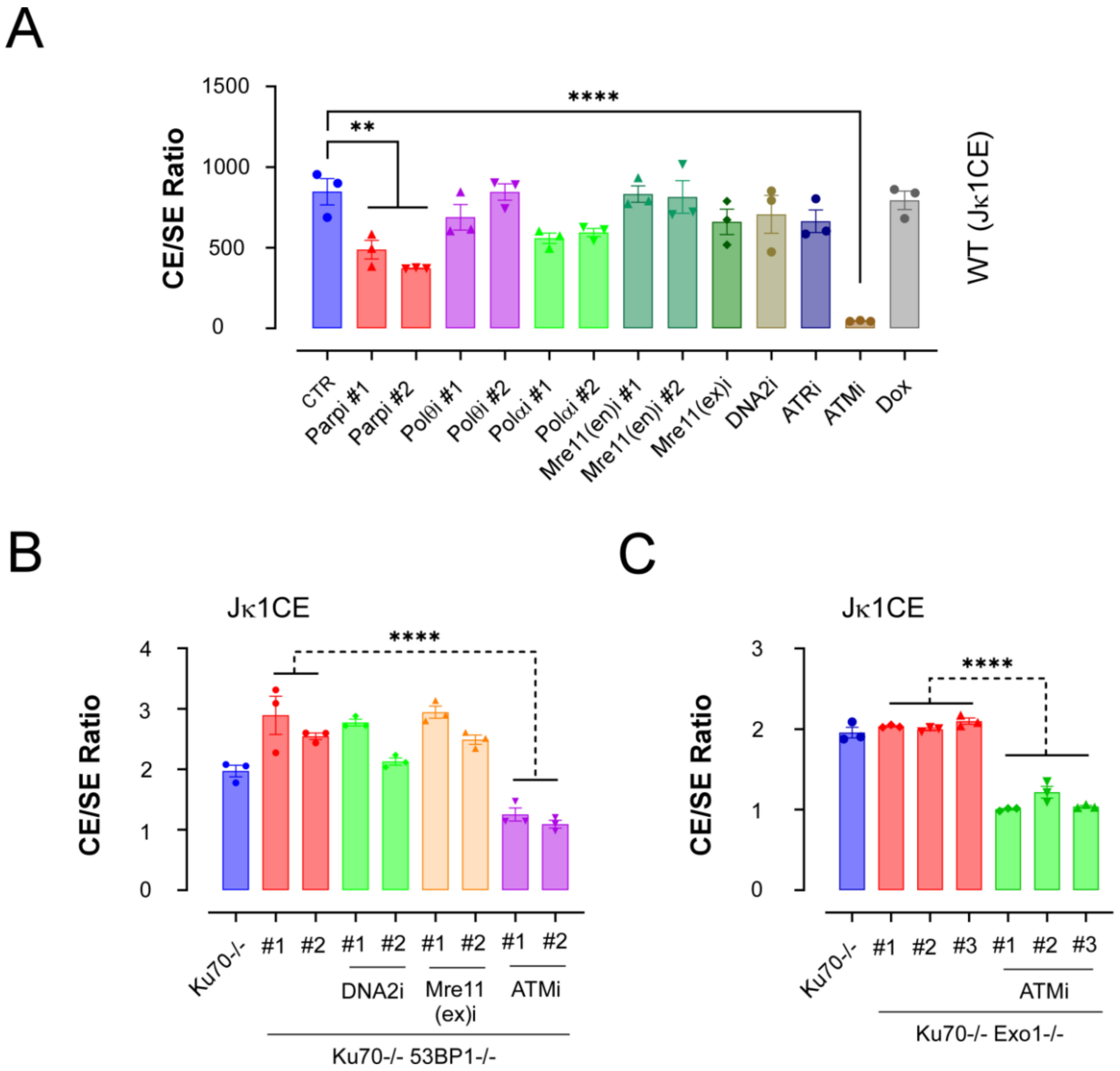


**Figure S16. ATM sustains the fidelity of coding end recombination.** (**A**) The coding to signal (CE/SE) ratios from the Jκ1CE bait are shown for WT with/without indicated inhibitors. (**B-C**) Same as (A) but *Ku70*$^{-/-}$ with/without *53bp1*$^{-/-}$ or *Exo1*$^{-/-}$ treated with/without indicated inhibitors. The difference between WT (CTR) and inhibitors are evaluated by One-way ANOVA plus post-comparison (A); differences for *Ku70*$^{-/-}$ *53bp1*$^{-/-}$ (B) and *Ku70*$^{-/-}$ *Exo1*$^{-/-}$ (C) are evaluated by Two-way ANOVA plus post-comparison, with ** ($p<0.01$) and **** ($p<0.0001$). All the experiments were independently repeated three times.

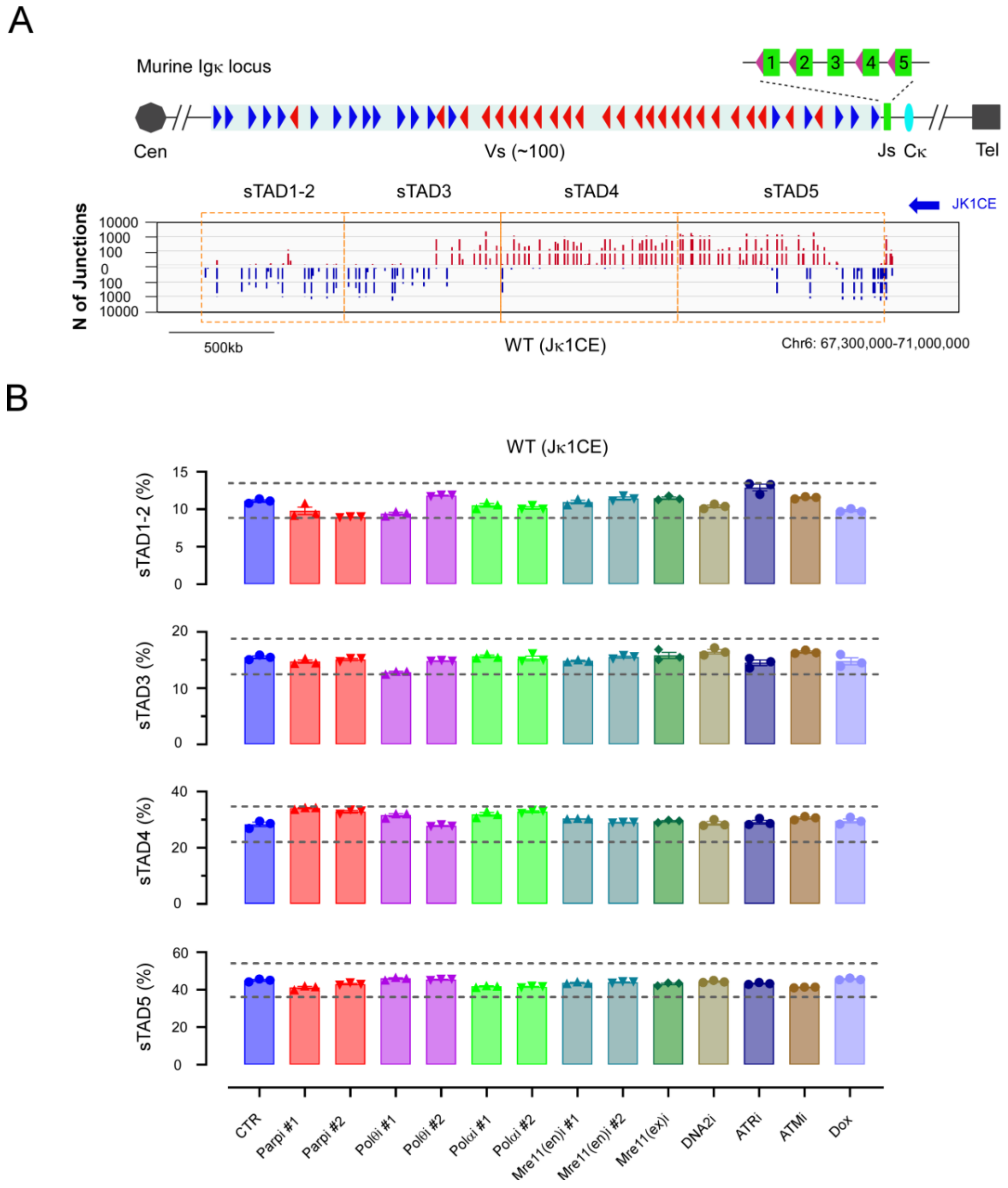


**Figure S17. The regional landscape of Vκ gene segment usage was not altered with treatment of the indicated compounds in WT cells.** (**A**) The Igκ locus is divided into four subTADs: sTAD1-2, sTAD3, sTAD4 and sTAD5, as shown in the *WT* control. (**B**) The percentage of sTAD1-2, sTAD3, sTAD4 and sTAD5 in *WT vAbl* cells with or without indicated compounds. The dash lines indicate the ± 20% threshold change of the CTR.

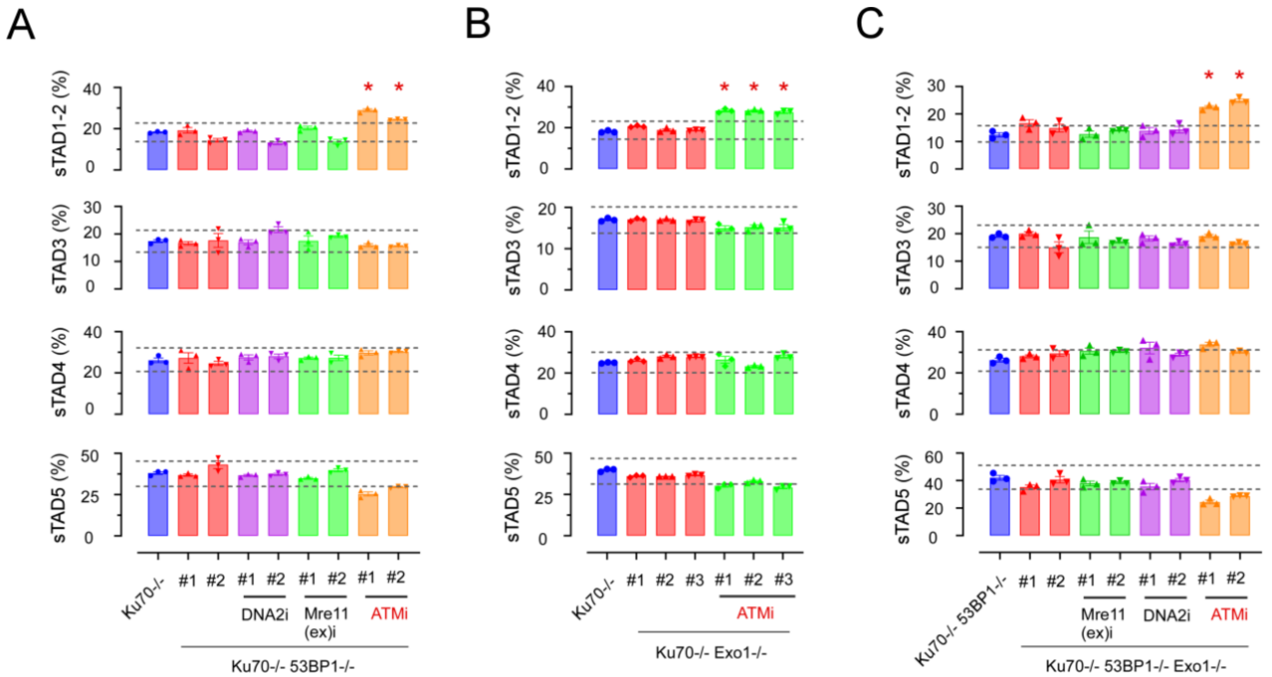


**Figure S18. The regional landscape of Vκ gene segment usage changes for additional A-EJ genetic backgrounds when ATM is inhibited.** (**A**) The Jκ1CE bait percentage of four sTADs (sTAD1-2, sTAD3, sTAD4 and sTAD5) in *vAbl Ku70*$^{-/-}$ cells and *Ku70*$^{-/-}$*53bp1*$^{-/-}$ with or without compounds including DNA2i, Mre11i and ATMi. The dash lines indicate the ± 20% threshold change of the CTR. (**B-C**) Same as (A) but for different treatment conditions as indicated.

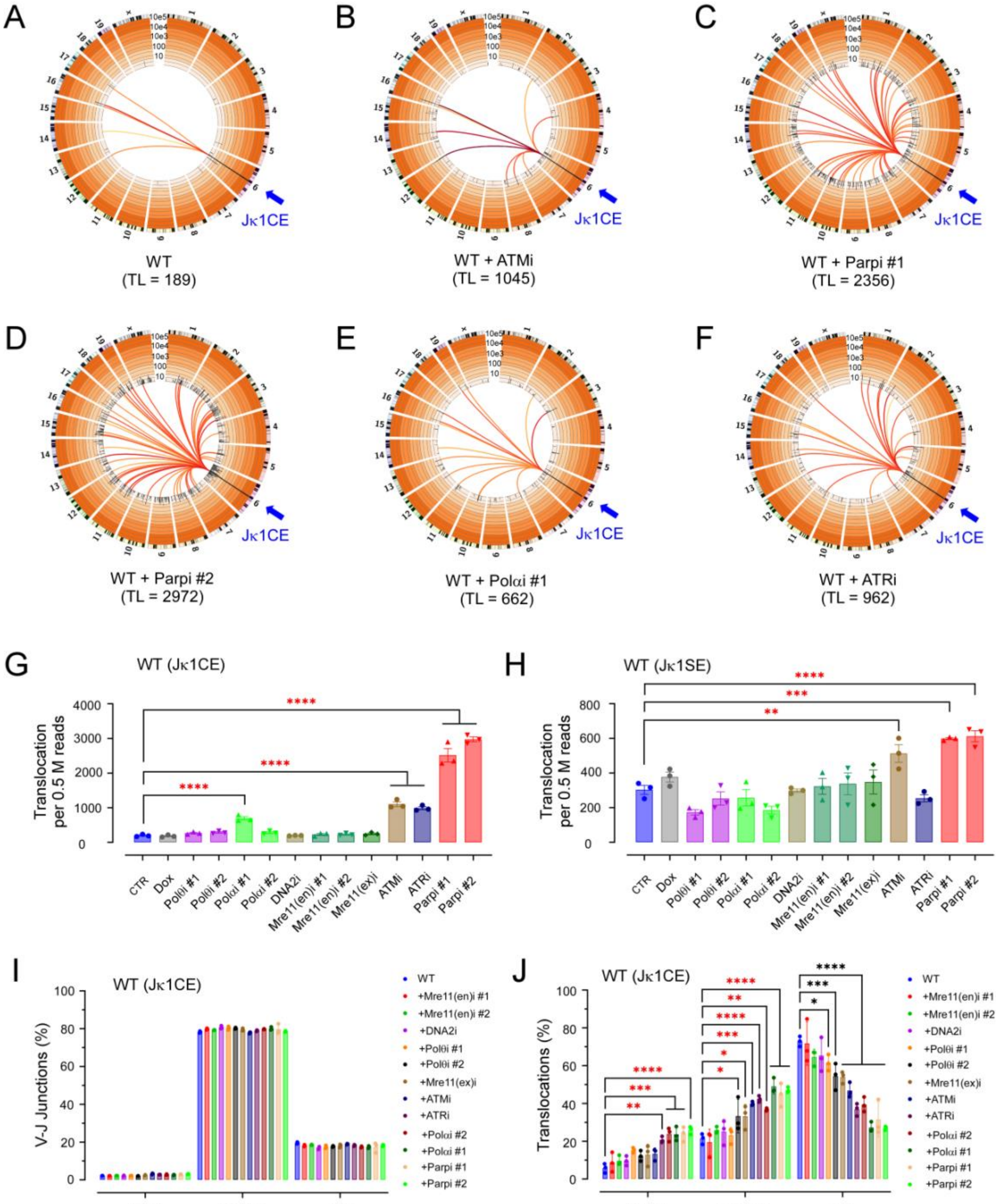


**Figure S19. Characterizing the genes that affect chromosome translocation in WT cells.** (**A-F**) The representative genome-wide plots of junctions joined with Jκ1CE in *vAbl WT* cells (A), +ATMi (B), +Parpi #1 (C), +Parpi #2 (D), +Polαi #1 (E) and +ATRi (F). (**G-H**) The relative translocation frequency per 0.5 M (million) reads in *vAbl* WT cells with/without indicated inhibitors captured by Jκ1CE (G) and Jκ1SE (H) baits, respectively. The numbers of translocation in *WT* (CTR) combined with indicated inhibitors were evaluated by one-way ANOVA plus post comparison with ** (p<0.01), *** (p<0.001) and **** (p<0.0001). (**I-J**) Junction structures of V region joints (I) and the absolute inter-chromosome translocations (J) in WT combined with indicated inhibitors. The junction structures were divided into three categories, microhomology larger than 1 bp (MH>1), direct repair plus the microhomology with 1 bp (Dir+MH1) and insertions (INS). The juncture structure of three categories between *WT* and the indicated inhibitors were evaluated by one-way ANOVA plus post comparison with * (p<0.05), ** (p<0.01), *** (p<0.001) and **** (p<0.0001). All experiments were independently repeated three times.

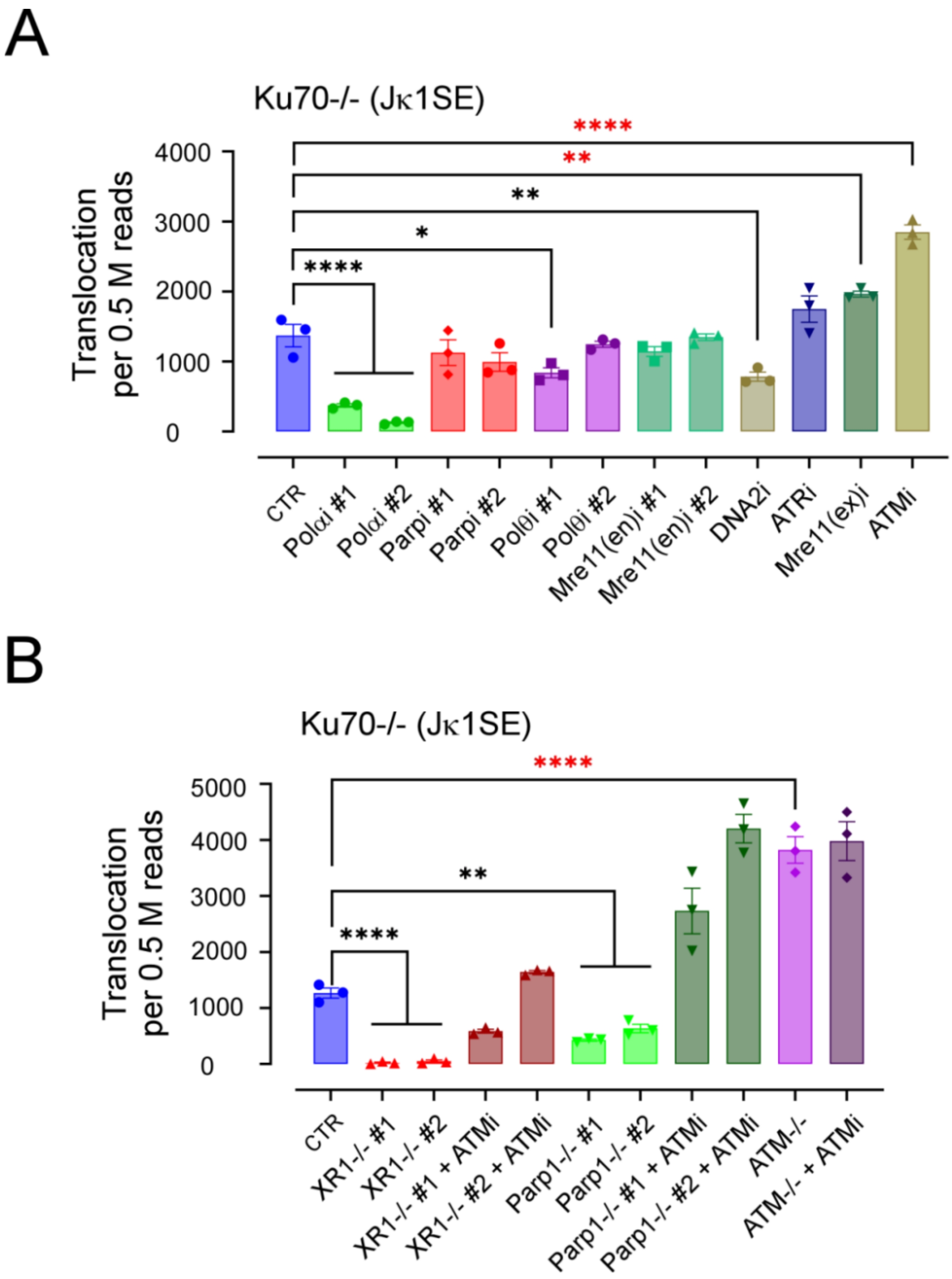


**Figure S20. Characterizing the genes that affect chromosome translocation of Jκ1SE in the absence of Ku70.** (**A**) The relative translocation frequency of Jκ1SE per 0.5 M (million) reads in *Ku70*$^{-/-}$ cells with/without indicated inhibitors. (**B**) Same as (A) but for XRCC1 (XR1), Parp1 and ATM deletion with/without ATMi. The numbers of translocation in *Ku70*$^{-/-}$ (CTR) combined with indicated inhibitors or gene modification or both were evaluated by one-way ANOVA plus post comparison with * ($p<0.05$), *** ($p<0.001$) and **** ($p<0.0001$). All experiments were independently repeated three times.

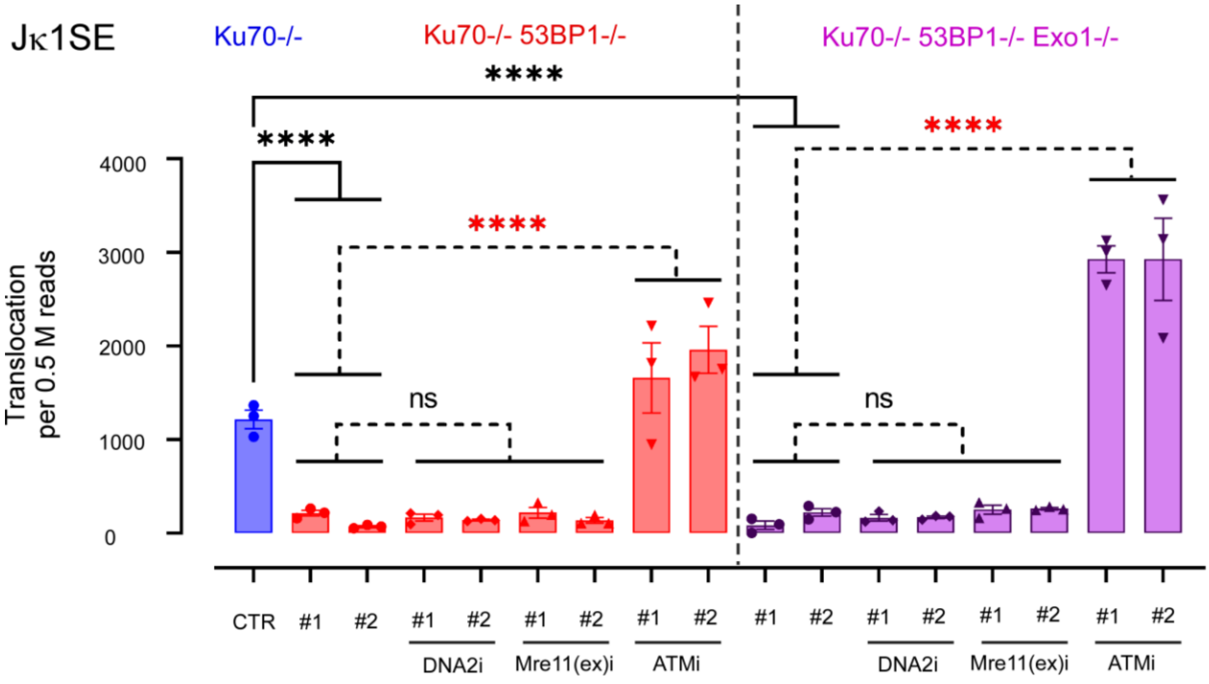


**Figure S21. 53BP1 and ATM counter each other, while Exo1 has little effect on chromosome translocation by A-EJ.** The relative translocation frequency from Jκ1SE bait per 0.5 M (million) reads in Ku70$^{-/-}$, *Ku70$^{-/-}$53bp1$^{-/-}$* and *Ku70$^{-/-}$53bp1$^{-/-}$Exo1$^{-/-}$* with/without indicated inhibitors. The numbers of translocation in *Ku70$^{-/-}$* (CTR) combined with indicated gene deletions were evaluated by one-way ANOVA plus post comparison; while that within *Ku70$^{-/-}$53bp1$^{-/-}$* (red bars) and *Ku70$^{-/-}$53bp1$^{-/-}$Exo1$^{-/-}$* (magenta bars) subgroups were evaluated by two-way ANOVA plus post comparison with **** ($p<0.0001$) and ns (no significance). All experiments were independently repeated three times.

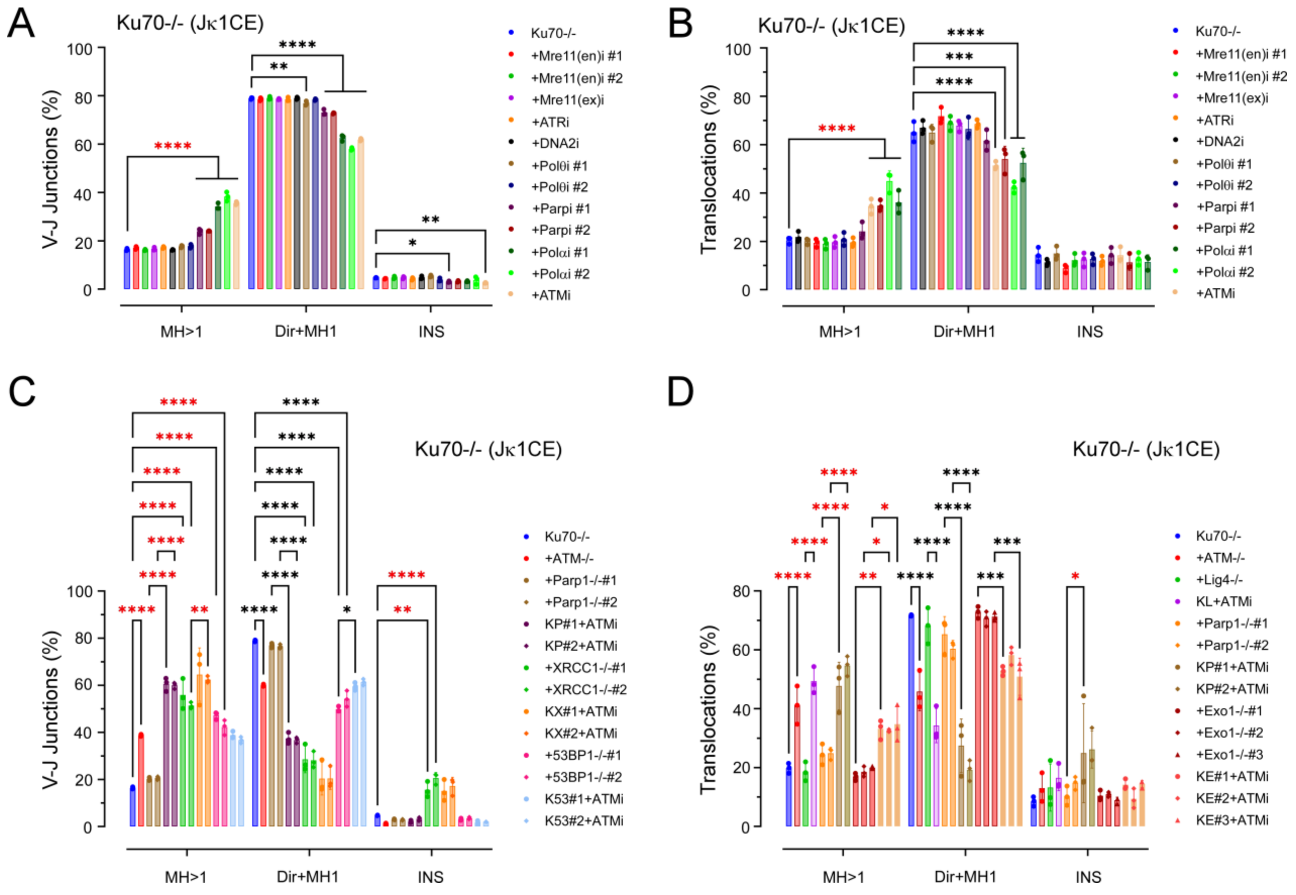


**Figure S22. The repair structure of V-J junctions and chromosomal translocations from Jκ1CE bait in *Ku70$^{-/-}$ vAbl* cells.** (**A-B**) Junction structures are divided into three categories: microhomology larger than 1 bp (MH>1), direct repair plus one microhomology (Dir+MH1) and insertions (INS). The proportion of these categories were measured for V-J junctions (A) and for absolute interchromosomal translocations (B) in *Ku70$^{-/-}$ vAbl* cells with/without inhibitors were quantified, respectively. (**C**) Same as (A) but for further gene deletions on top of *Ku70$^{-/-}$*: *Ku70$^{-/-}$Atm$^{-/-}$* (*+ATM$^{-/-}$*), *Ku70$^{-/-}$Parp1$^{-/-}$* (*+Parp1$^{-/-}$* #1/2), *Ku70$^{-/-}$Parp1$^{-/-}$* with ATMi (*KP* #1/2 +ATMi), *Ku70$^{-/-}$Xrcc1$^{-/-}$* (*+XRCC1$^{-/-}$* #1/2), *Ku70$^{-/-}$Xrcc1$^{-/-}$* with ATMi (*KX* #1/2 + ATMi), *Ku70$^{-/-}$53bp1$^{-/-}$* (*+53BP1$^{-/-}$* #1/2) and *Ku70$^{-/-}$53bp1$^{-/-}$* with ATMi (*K53* #1/2 +ATMi). (**D**) Same as (B) but for further gene deletions on top of *Ku70$^{-/-}$*: *Ku70$^{-/-}$Atm$^{-/-}$* (*+ATM$^{-/-}$*), *Ku70$^{-/-}$Lig4$^{-/-}$* (*+Lig4$^{-/-}$*), *Ku70$^{-/-}$Lig4$^{-/-}$* with ATMi (*KL* + ATMi), *Ku70$^{-/-}$Parp1$^{-/-}$* (*+Parp1$^{-/-}$* #1/2), *Ku70$^{-/-}$Parp1$^{-/-}$* with ATMi (*KP* #1/2 +ATMi), *Ku70$^{-/-}$Exo1$^{-/-}$* (*+Exo1$^{-/-}$* #1/2/3) and *Ku70$^{-/-}$Exo1$^{-/-}$* with ATMi (*KE* #1/2/3 +ATMi). The difference among each group was evaluated by one-way ANOVA plus post comparison with * ($p<0.05$), ** ($p<0.01$), *** ($p<0.001$) and **** ($p<0.0001$). All the experiments were independently repeated three times.
